# Endogenous GDF15 and FGF21 additively alleviate hepatic steatosis and insulin resistance in obese mice

**DOI:** 10.1101/2022.06.08.495255

**Authors:** Satish Patel, Afreen Haider, Anna Alvarez-Guaita, Guillaume Bidault, Julia Sarah El-sayed Moustafa, Esther Guiu-Jurado, John A. Tadross, James Warner, James Harrison, Samuel Virtue, Fabio Scurria, Ilona Zvetkova, Matthias Blüher, Kerrin S. Small, Stephen O’Rahilly, David B. Savage

## Abstract

Obesity in mice and humans is associated with elevated levels of at least two hormones responsive to cellular stress, namely GDF15 and FGF21. Over-expression of each of these is associated with weight loss and beneficial metabolic changes but where they are secreted from and what they are required for physiologically in the context of overfeeding remains unclear. Here we used tissue selective knockout mouse models to establish that, like FGF21, circulating GDF15 is primarily derived from the liver, rather than adipose tissue, muscle or macrophages in high fat fed mice. Combined whole body deletion of FGF21 and GDF15 does not result in any additional weight gain in high fat fed mice but is associated with significantly greater hepatic steatosis and insulin resistance. Collectively the data suggest that activation of the integrated stress response in hepatocytes is a major driver for GDF15 and FGF21 secretion in the context of overfeeding, and that they both act to alleviate this metabolic stress.

## Introduction

Healthy adipose tissue is essential when coping with sustained nutritional overload and manifests a remarkable capacity to increase in size (Arner and Rydén, 2021). Depending on factors such as the duration and macronutrient source of excess caloric intake, fat mass can double in humans (Tremblay et al., 1992) and quadruple in rodent models (Carpenter et al., 2013). Ultimately however, adipocytes start to die triggering a chronic inflammatory response, in which macrophages play a key role (Cinti, 2012; Lindhorst et al., 2021; Murano et al., 2008; Skurk et al., 2007; Virtue and Vidal-Puig, 2010; Weisberg et al., 2003). Since excess energy cannot normally be excreted and humans, at least, manifest very small increases in energy expenditure when overfed (Savage et al., 2005), it necessarily accumulates in other ectopic sites (and in plasma lipoproteins to some extent), setting off a cascade of metabolic complications. In the liver this ectopic lipid accumulation is strongly associated with the development of insulin resistance (Kim et al., 2001; Korenblat et al., 2008; Loomba et al., 2021; Marchesini et al., 1999; Roden and Shulman, 2019; Samuel and Shulman, 2018; Sanyal et al., 2001) and in some cases a hepatic inflammatory response. As many as a third of patients with non-alcoholic fatty liver disease (NAFLD) progress to cirrhosis and some of these go on to develop hepatocellular carcinoma (Huang et al., 2021; Loomba et al., 2020). Lipid also accumulates in skeletal muscle (Kim et al., 2001; Krssak et al., 1999; Pan et al., 1997) where it has again been linked to the pathogenesis of insulin resistance and type 2 diabetes (Petersen and Shulman, 2017; Roden and Shulman, 2019). Although there are some reports of ‘inflammation’ in skeletal muscle in this context, the underlying mechanisms, and the impact on muscle itself and on systemic insulin resistance are less clear (Wu and Ballantyne, 2017).

Ectopic lipid accumulation in the liver has been reported by several groups to be associated with activation of the integrated stress response (ISR) (Gregor et al., 2009; Hotamisligil, 2010; Nakatani et al., 2005; Ozcan et al., 2004), whereas in skeletal muscle of humans or high-fat fed mice there is limited evidence of induction of the ISR (Deldicque et al., 2011; Koh et al., 2013; Ozcan et al., 2004). The ISR is triggered by a variety of stressors (such as endoplasmic reticulum stress, mitochondrial stress, hypoxia and, amino acid or glucose depletion) which activate one or more of at least four kinases leading to the phosphorylation of eukaryotic initiation factor 2 alpha (eIF2α) (Costa-Mattioli and Walter, 2020). This in turn attenuates overall protein synthesis whilst permitting selective translation of specific proteins required for cellular adaptation, repair and alteration of metabolic homeostasis (Hotamisligil and Davis, 2016; Pakos-Zebrucka et al., 2016). We (Patel et al., 2019) and others (Chung et al., 2017; Dong et al., 2015; Laeger et al., 2014) have shown that activation of the ISR (via nutritional, genetic or pharmacological stressors) is associated with increased expression of growth differentiation factor 15 (GDF15) and fibroblast growth factor 21 (FGF21). Plasma levels of both FGF21 (Gómez-Ambrosi et al., 2017; Kharitonenkov et al., 2005; Kralisch et al., 2013; Reinehr et al., 2012; Zhang et al., 2008) and GDF15 (Carballo-Casla et al., 2021; Vila et al., 2011; Xiong et al., 2017) are known to be increased in obese humans and rodents as well as in other metabolic disease states such as insulin resistance (Chavez et al., 2009; Kempf et al., 2012), NAFLD (Bilson et al., 2021; Dushay et al., 2010; Galuppo et al., 2022; Kim et al., 2018; Rusli et al., 2016; Tucker et al., 2019) and mitochondrial disease (Poulsen et al., 2020; Suomalainen et al., 2011). Both stress-induced cytokines, GDF15 and FGF21, have attracted considerable interest as potential therapies for obesity and its associated metabolic disease (Keipert and Ost, 2021).

FGF21 was discovered in 2000 and reported to be primarily expressed in the liver (Nishimura et al., 2000) though it is now known to be more widely expressed, including other key metabolic tissues such as adipose tissue, skeletal muscle and pancreas (Fisher and Maratos-Flier, 2016; Kliewer and Mangelsdorf, 2019). FGF21 was found to be a potent regulator of glucose uptake in an *in vitro* screen in 3T3-L1 adipocytes (Kharitonenkov et al., 2005) and has subsequently been associated with a range of pleiotropic effects, including improved insulin sensitivity and β cell function, reduced hepatic lipogenesis, and increased energy expenditure via brown fat thermogenesis (Flippo and Potthoff, 2021). Using tissue-specific knockout mice, plasma FGF21 was shown to be primarily derived from the liver in response to high-fat feeding (Markan et al., 2014); it then acts centrally to regulate energy expenditure and body weight (Douris et al., 2015; Owen et al., 2014). FGF21 is also reported to act in an auto/paracrine fashion when secreted by adipocytes and pancreatic exocrine cells (Coate et al., 2017; Dutchak et al., 2012; Fisher et al., 2012; Singhal et al., 2016). Molecularly, FGF21 signals via the FGF receptor-1c and its cognate co-receptor β-klotho which is predominantly expressed in target tissue such as the CNS and adipocytes (Adams et al., 2012; Ding et al., 2012; Foltz et al., 2012). Both transgenic overexpression and exogenous administration of supraphysiologic levels of FGF21 in genetic- or diet-induced obese (DIO) rodents, substantially reduces body weight, hypertriglyceridaemia and hyperglycaemia (Coskun et al., 2008; Kharitonenkov et al., 2005; Wente et al., 2006). These responses were shown to be associated with reductions in hepatic and intramyocellular lipid content (TAG and DAG)(Camporez et al., 2013). Beneficial metabolic effects of exogenous FGF21 administration were also observed in obese or diabetic primates (Kharitonenkov et al., 2007; Talukdar et al., 2016) and in humans (Gaich et al., 2013; Talukdar et al., 2016), although the improvements in hyperglycaemia were disappointingly modest in humans. FGF21 deficiency in mice has been linked with reduced insulin sensitivity in a high-fat diet setting (Li et al. 2018), but the impact on body weight remains unclear as studies have reported both lower and higher body weight in FGF21 null mice (Adams et al., 2013; Li et al., 2018). Some studies suggested that the weight of FGF21 knockout mice was increased early after transition to a high fat diet but that this difference was subsequently lost (Fisher et al., 2010; Singhal et al., 2016).

GDF15, originally identified as a gene upregulated in activated macrophages (Bootcov et al., 1997), can be expressed in almost all cell types, and is relatively highly expressed in several tissues including the liver, kidneys, intestines and especially the placenta. Plasma GDF15 is elevated in a range of human diseases, in addition to obesity and the metabolic syndrome, where it is widely considered to be a useful biomarker (Breit et al., 2021; Lockhart et al., 2020; Wischhusen et al., 2020). GDF15 mRNA expression is elevated within the liver and adipose tissue of high-fat fed mice (Patel et al., 2019). GDF15 null mice weigh more than WT littermates on a high fat diet and are glucose intolerant (Tran et al., 2018). Meanwhile, GDF15 over-expressing transgenic mice are protected from diet-induced obesity and display improved insulin sensitivity (Macia et al., 2012). Concordantly, pharmacological treatment with recombinant GDF15 reduces body weight and food intake in obese rodents and primates (Chrysovergis et al., 2014; Johnen et al., 2007; Xiong et al., 2017). These metabolic impacts of GDF15 are mediated via the GFRAL receptor along with its tyrosine kinase coreceptor Ret in the hindbrain (Emmerson et al., 2017; Mullican et al., 2017).

Here we sought firstly to clarify the principal source of GDF15 in HFD fed mice. Having shown that the liver is a major source of GDF15 in this context, similarly to what has previously been reported for FGF21, we proceeded to evaluate the phenotypic impact of deleting both genes in mice. As both are present at least as far back as zebrafish (Itoh, 2007; Pereiro et al., 2020) and both have been implicated in weight loss, we hypothesized that they might act synergistically to alleviate the stress imposed by nutritional overload so we generated and characterised GDF15:FGF21 double knockout mice.

## Results

### Impact of GDF15 deletion on body weight regulation in mice

In order to confirm prior reports suggesting that GDF15 null mice are heavier when fed a high-fat diet (HFD) (Tran et al., 2018), wild-type (WT) and GDF15 KO mice were fed a 60% HFD for ∼20 weeks. The GDF15 KO mice displayed a small but significant increase in absolute body weight compared to their WT littermates **(Figure 1A)**, the difference becoming evident from about 10 weeks into the HFD, particularly when the data was displayed as % body weight gain **(Figure 1B)**. Subcutaneous, epididymal and brown adipose tissue weights were similar between the groups, but GDF15 KO mice manifested increased liver tissue mass **(Figure 1C)**. Biochemical **(Figure 1D)** and histological analyses revealed elevated hepatic lipid content in GDF15 KO mice **(Figure 1E and Figure 1 supplemental)** and plasma triglycerides; cholesterol, leptin and liver enzyme concentrations were also higher than those of wildtype littermates **(Figure 1F)**. Blood glucose concentrations were similar in 6 hour fasted mice whereas plasma insulin and HOMA-IR were elevated suggesting mild insulin resistance **(Figure 1G)**. In keeping with these data, glucose and insulin tolerance were mildly impaired in the GDF15 KO group **(Figure 1H-I).** High-fat feeding is known to increase plasma FGF21 in mice (Patel et al., 2019) and was even higher in GDF15 KO mice after 16 weeks of HFD feeding **(Figure 1J)**. In keeping with this observation, FGF21 mRNA was increased in the liver of the GDF15 KOs but not in WAT, BAT or skeletal muscle **(Figure 1K)**.

**Figure 1:**
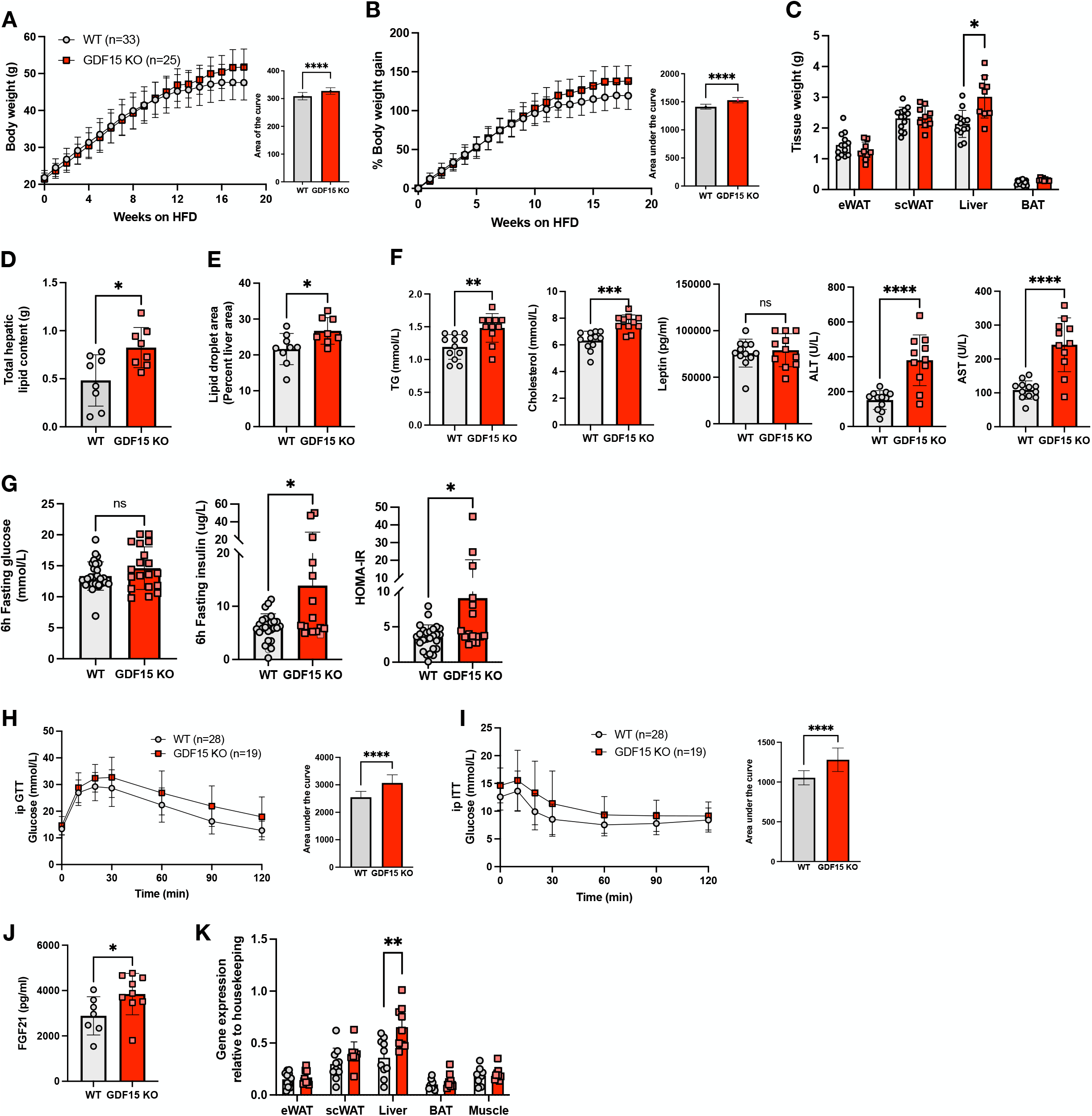
Phenotypic characterization of GDF15 knockout mice on a high-fat diet (HFD) (A and B) Body weight and percent body weight gain of wild type (WT) and GDF15 KO mice fed a 60% HFD; Inset, area of the curve analysis of body weight over time and area under the curve analysis of % body weight gain over time. (C) Weight of epididymal white adipose tissue (eWAT), subcutaneous white adipose tissue (scWAT), liver and brown adipose tissue (BAT), harvested at the end of the study, 25 weeks of HFD (n=13,9). (D) Weight of total lipids isolated per mg of liver, normalized to total liver weight (n=8). (E) Lipid droplet area (Percent liver area) determined from histological analyses of haematoxylin/eosin (H&E) stained liver sections (n=9,8) (F) Plasma triglyceride (TG), cholesterol, leptin, alanine transaminase (ALT) and aspartate transaminase (AST) from random fed mice after 16 weeks of HFD-feeding (n=12,11) (G) Blood glucose, plasma insulin and HOMA-IR levels from 6h fasted mice, after 16 weeks of HFD feeding (n=16-29) (H and I) Blood glucose levels during intraperitoneal (ip) glucose tolerance test (GTT) and insulin tolerance test (ITT) after 16 weeks of HFD feeding. Inset, area under the curve analysis of glucose over time. (J) Plasma FGF21 levels from random fed mice at 16 weeks of HFD feeding (n=7,9) (K) Fgf21 mRNA expression in tissues from WT and GDF15 KO mice after 25 weeks HFD feeding (n=8-11) All data are means ± S.D */**/***/**** - p<0.05/0.01/0.001/0.0001

### GDF15 is upregulated in adipose tissue macrophages and hepatocytes

Having previously shown that high fat feeding in mice is associated with elevated plasma GDF15 levels and increased GDF15 mRNA in white and brown adipose tissue, and the liver, we used RNA scope to identify the cellular source. In white adipose tissue, the vast majority of GDF15 expression was detected in macrophages **(Figure 2A).** In chow fed mice, RT-PCR quantification confirmed that GDF15 mRNA was mainly expressed in myeloid cells (CD11b+ SVF) with very little expression in adipocytes or CD11b-SVF (stromal vascular fraction) **(Figure 2B)**. In HFD fed mice, GDF15 was readily detectable within all three eWAT fractions. However, GDF15 expression within the CD11b-SVF and adipocyte fractions is likely due to macrophage contamination, as they express the macrophage marker EMR1 which correlates with GDF15 expression **(Figure 2C-D)**.

**Figure 2:**
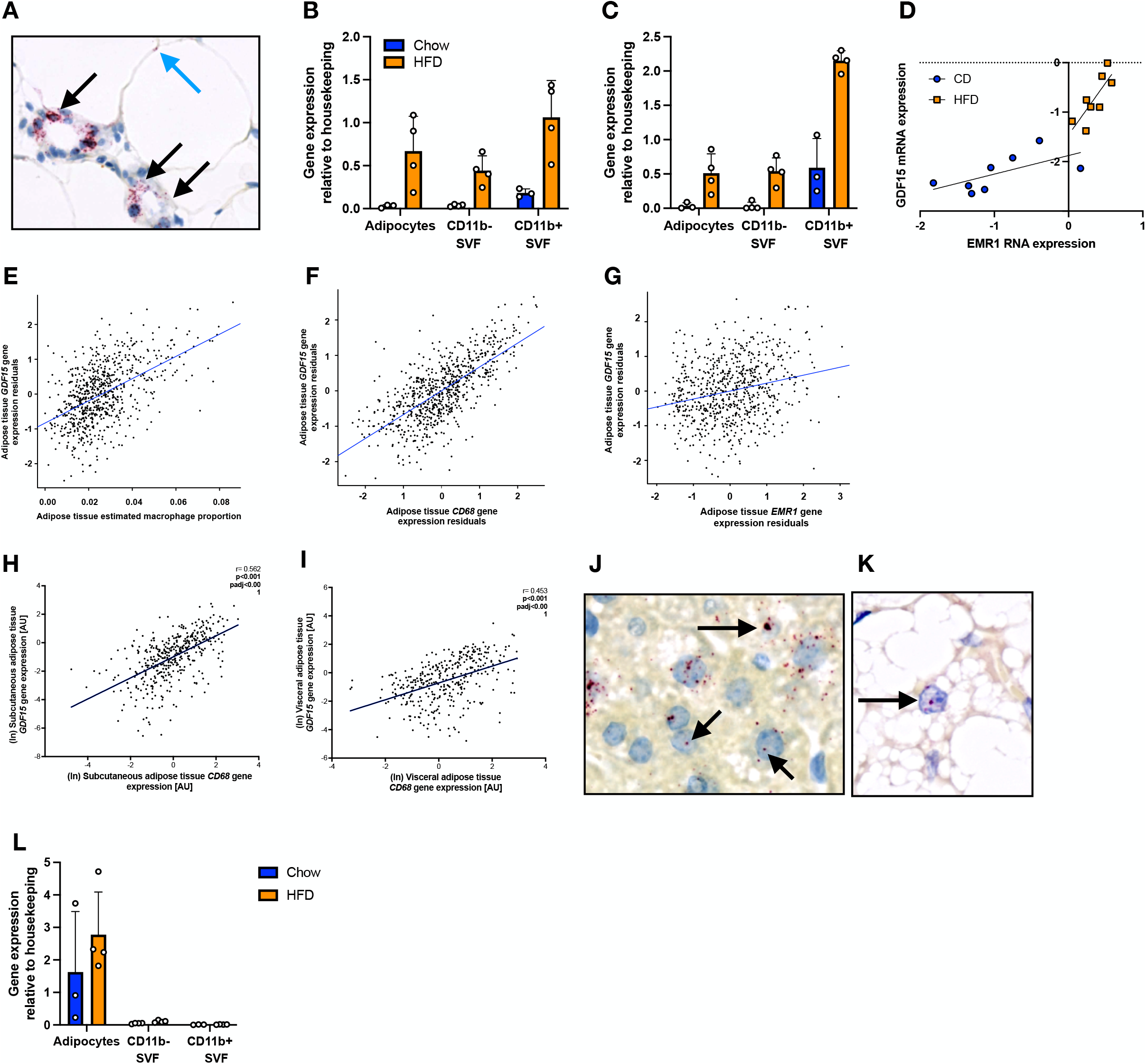
Gdf15 mRNA expression within high-fat diet-fed mouse and human adipose tissue. (A) In situ hybridization analysis of Gdf15 mRNA (red) from 18 week old high fat diet (HFD) fed wild type (WT) mouse epididymal adipose tissue and liver tissue. Black arrows indicate Gdf15 staining in foamy macrophages. Blue arrow indicates Gdf15 staining within adipocytes. (B and C) Gdf15 and EMR1 mRNA expression from 14 week old Chow or HFD fed WT mouse epididymal tissue fractionated into adipocytes, CD11b negative (-) and CD11b positive (+) stromal vascular fractions (SVF) (n=3-4). (D) Correlation of GDF15 expression with EMR1 expression in epididymal tissue from 14 week Chow or HFD fed wild type mice (n=8) (E-G) Human subcutaneous adipose tissue GDF15 gene expression levels in the TwinsUK adipose study associated with estimated macrophage proportion in adipose tissue (E), and macrophage markers CD68 and EMR1 (F and G). Each point represents data from a single individual. Plotted gene expression residuals of GDF15, CD68 and EMR1 were adjusted for age, BMI and RNA-Seq technical covariates. (H and I) GDF15 gene expression levels in the ‘Obese study’ associated with macrophage marker CD68 in human subcutaneous and visceral adipose tissue. (J and K) In situ hybridization analysis of Gdf15 mRNA (red) from 18 week old HFD fed WT mouse liver and brown adipose tissue. (L) Fgf21 mRNA expression from 14 week Chow or HFD fed WT mouse epididymal tissue fractionated in to adipocytes, CD11b negative (-) and CD11b positive (+) stromal vascular fraction (SVF) (n=3-4).

In keeping with these murine data, GDF15 mRNA expression in human subcutaneous adipose tissue samples in 733 individuals from the TwinsUK cohort was strongly associated with the estimated adipose tissue macrophage proportion (β=32.81; *P* < 2.2×10^-16^) (**Figure 2E)**, as well as with expression of macrophage markers CD68 (β=0.68; *P* < 2.2×10^-16^) and EMR1 (β=0.21; *P*=3.5×10^-9^) **(Figure 2F-G)**. We confirmed these findings in a second large obese cohort (n=525) in which we also had access to paired visceral adipose tissue samples. In this sample set, GDF15 mRNA expression was again strongly correlated with CD68 mRNA in subcutaneous fat. The correlation was also present in visceral fat, though interestingly was slightly weaker in that depot (Subcutaneous fat: r=0.562, *P* < 0.001; Visceral fat: r=0.453, *P* < 0.001). Both associations withstand adjustment for age, sex and BMI (*Padj* < 0.001) (**Figure 2H-I).**

In liver samples from HFD fed mice, the RNA scope analysis suggested that GDF15 mRNA was primarily detectable in hepatocytes with no visually apparent contribution from non-hepatocyte cell populations **(Figure 2J)**. Previously published work has suggested that GDF15 expression is also increased in human liver tissue samples in the context of NAFLD (Govaere et al., 2020). In brown adipose tissue, the GDF15 mRNA signal appeared to originate predominantly from selected brown adipocytes **(Figure 2K).** In contrast, HFD-induced FGF21 is reported to be predominantly expressed within adipocytes themselves whereas within the liver it is also expressed in hepatocytes (Markan et al., 2014). We confirmed the former using RT-PCR in epididymal WAT fractions **(Figure 2L)**.

### Characterisation of macrophage specific GDF15 KO mouse models

As macrophages appear to be the predominant site of GDF15 expression in adipose tissue we proceeded to generate myeloid-specific GDF15 knockout mice (KO) using Cre recombinase under the control of the Lysozyme M promoter (LysM Cre). LysM-GDF15^KO^ were viable, fertile and born with no obvious abnormalities. Deletion of GDF15 was confirmed within isolated bone marrow derived macrophages (BMDMs), where we observed a >90% reduction in GDF15 mRNA **(Figure 3 supplemental)**. In keeping with these data, secretion of GDF15 into the media from isolated GDF15 null BMDMs was undetectable **(Figure 3 supplemental)**. To verify that GDF15 was effectively deleted within the myeloid population *in vivo*, adipose tissue macrophages (ATMs) were purified from WT and LysM-GDF15^KO^ epididymal adipose tissue. Like in BMDMs, mRNA analysis revealed >90% reduction of GDF15 expression in the LysM-GDF15^KO^ ATMs **(Figure 3A)**. Furthermore, analysis of gene expression from whole epididymal WAT displayed significant loss (>90%) of GDF15 expression in LysM-GDF15^KO^ mice but not in other tissues **(Figure 3B)**.

**Figure 3:**
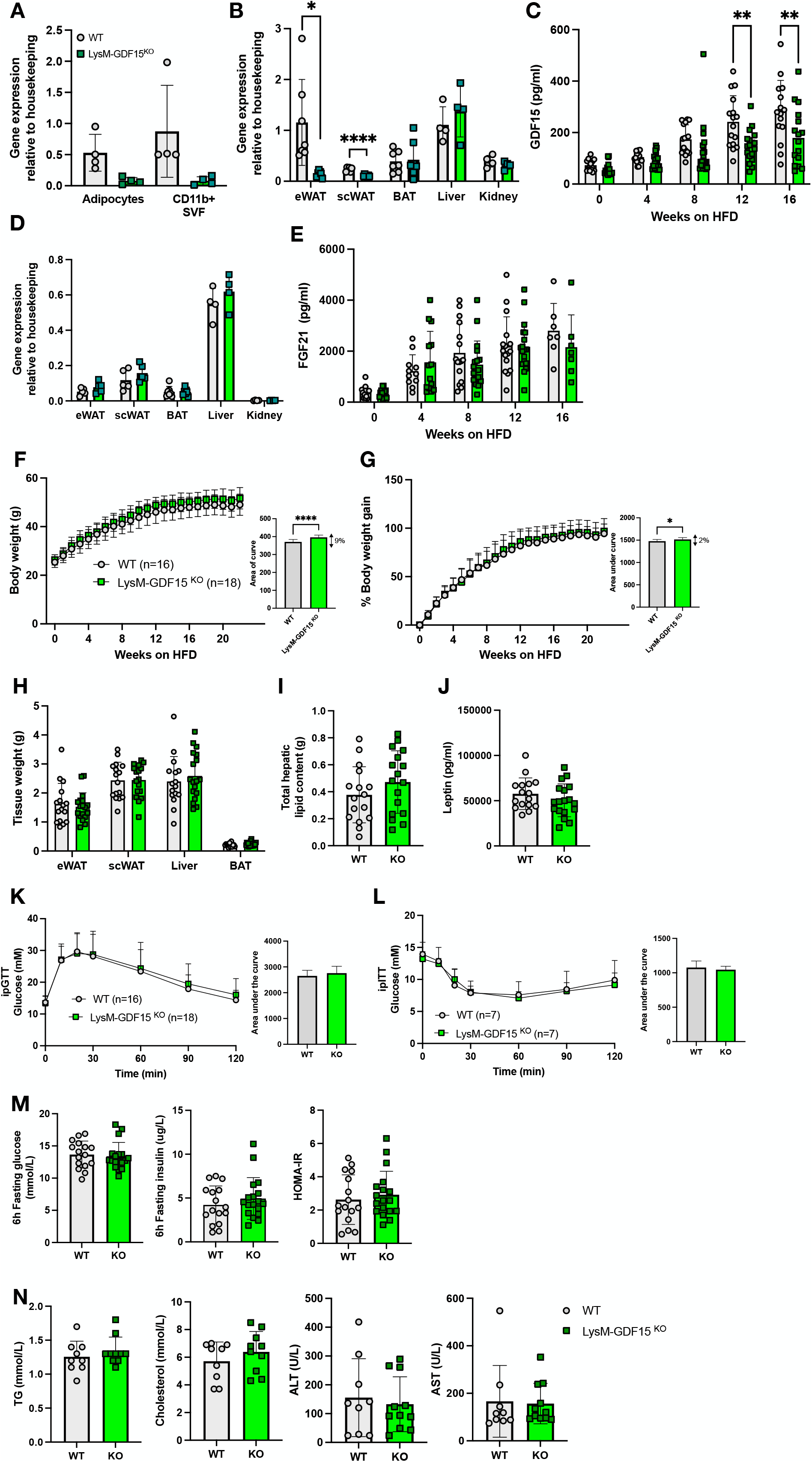
Phenotypic characterization of LysM-Cre mediated GDF15 macrophage knockout mouse on a high fat diet (HFD) (A) Gdf15 mRNA expression from 24 week old high fat diet (HFD) fed wild type (WT) and LysM-GDF15^KO^ mouse epididymal adipose tissue fractionated into adipocytes and CD11b positive (+) stromal vascular fraction (SVF) (n=4). (B) Gdf15 mRNA expression in tissues from 24 week HFD fed WT and LysM-GDF15^KO^ mice. Epididymal white adipose tissue (eWAT), subcutaneous white adipose tissue (scWAT), and brown adipose tissue (BAT) (n=4-7). (C) Plasma GDF15 levels from random fed mice at indicated weeks on HFD (n=15-18) (D) Tissue Fgf21 mRNA expression in tissues from 24 week HFD fed WT and LysM-GDF15^KO^ mice (n=4-7). (E) Plasma FGF21 levels from random fed mice at indicated weeks on HFD (n=11-18). (F and G) Body weight and percent body weight gain of WT and LysM-GDF15^KO^ mice fed a 60% HFD; Inset, area of curve analysis of body weight over time and area under the curve analysis of % body weight gain over time. (H) Weight of epididymal white adipose tissue (eWAT), subcutaneous white adipose tissue (scWAT), liver and brown adipose tissue (BAT) harvested at the end of the study, 24 weeks of HFD (n=16-18). (I) Weight of total lipids isolated per mg of liver, normalized to total liver weight (n=16,17). (J) Plasma leptin levels in mice from random fed mice after 16 weeks HFD-feeding (n=16-18). (K and L) Blood glucose levels during ip glucose tolerance test (GTT) and insulin tolerance test (ITT) after 16 weeks of HFD feeding. Inset, area under the curve analysis of glucose over time. (M) Blood glucose, plasma insulin and HOMA-IR levels from 6 h fasted mice, after 16 weeks of HFD feeding (n=16,18). (N) Plasma triglyceride (TG), cholesterol, alanine transaminase (ALT) and aspartate transaminase (AST) from random fed mice, after 16 weeks of HFD feeding (n=9-11). All data are means ± S.D */** - p<0.05/0.01

As expected, plasma concentrations of GDF15 increased during HFD feeding in all mice, but GDF15 concentrations were ∼30% lower in the LysM-GDF15^KO^ cohort **(Figure 3C)**. FGF21 gene expression and plasma concentrations were similar between genotypes **(Figure 3D,E)**. Weekly weight analysis revealed that there was a trend for LysM-GDF15^KO^ mice to be slightly heavier on a HFD compared to their WT counterparts **(Figure 3F-G)** though tissue weights, hepatic lipid content and plasma leptin were similar to WT mice **(Figure 3H-J)**. Glucose and insulin tolerance were also similar to that of WT littermates **(Figure 3K-L)** along with glucose and insulin concentrations **(Figure 3M).** Plasma lipids (TG and cholesterol) were similar, as were plasma liver enzyme concentrations **(Figure 3N)**.

Since the LysM-Cre driven deletion is present from birth and specifically targets the myeloid population, it is possible that compensatory mechanisms might account for the very modest impact on plasma GDF15 concentrations. To test this possibility, we transplanted bone marrow from GDF15 KO mice to regenerate the hematopoietic lineage in irradiated WT mice. GDF15 mRNA expression was effectively deleted within the BMT-GDF15^KO^ WAT depots but not in other tissues (liver and BAT) **(Figure 4A),** but the plasma GDF15 concentrations were similar in both lines **(Figure 4B)**. It’s worth noting that the GDF15 concentrations are substantially higher in the BMT groups **(Figure 4B)** than in the LysM-Cre cohorts **(Figure 3C)**. We presume that this relates to the irradiation therapy which is known to increase GDF15 (Moritake et al., 2012; Okazaki et al., 2006). However, similarly to what was observed in the whole body GDF15 null mice, plasma FGF21 concentrations and hepatic FGF21 mRNA expression were increased in the BMT-GDF15^KO^ mice **(Figure 4C-D)**. When both groups were fed a 60% HFD, the BMT-GDF15^KO^ cohort displayed slightly greater weight gain than the WT **(Figure 4E)**. This difference became evident 8 weeks after HFD feeding, particularly when the data was analysed as % body weight gain **(Figure 4F)**. Tissue weight analyses revealed a slight reduction in epididymal fat mass and a trend for increased liver weight **(Figure 4G)** but plasma TG, cholesterol, leptin concentrations as well as liver enzymes were similar **(Figure 4H).** Furthermore, there was no difference in hepatic fat content **(Figure 4I)**. In terms of glucose tolerance, insulin sensitivity, fasting glucose and insulin levels, both sets of animals had similar profiles **(Figure 4J-L)**.

**Figure 4:**
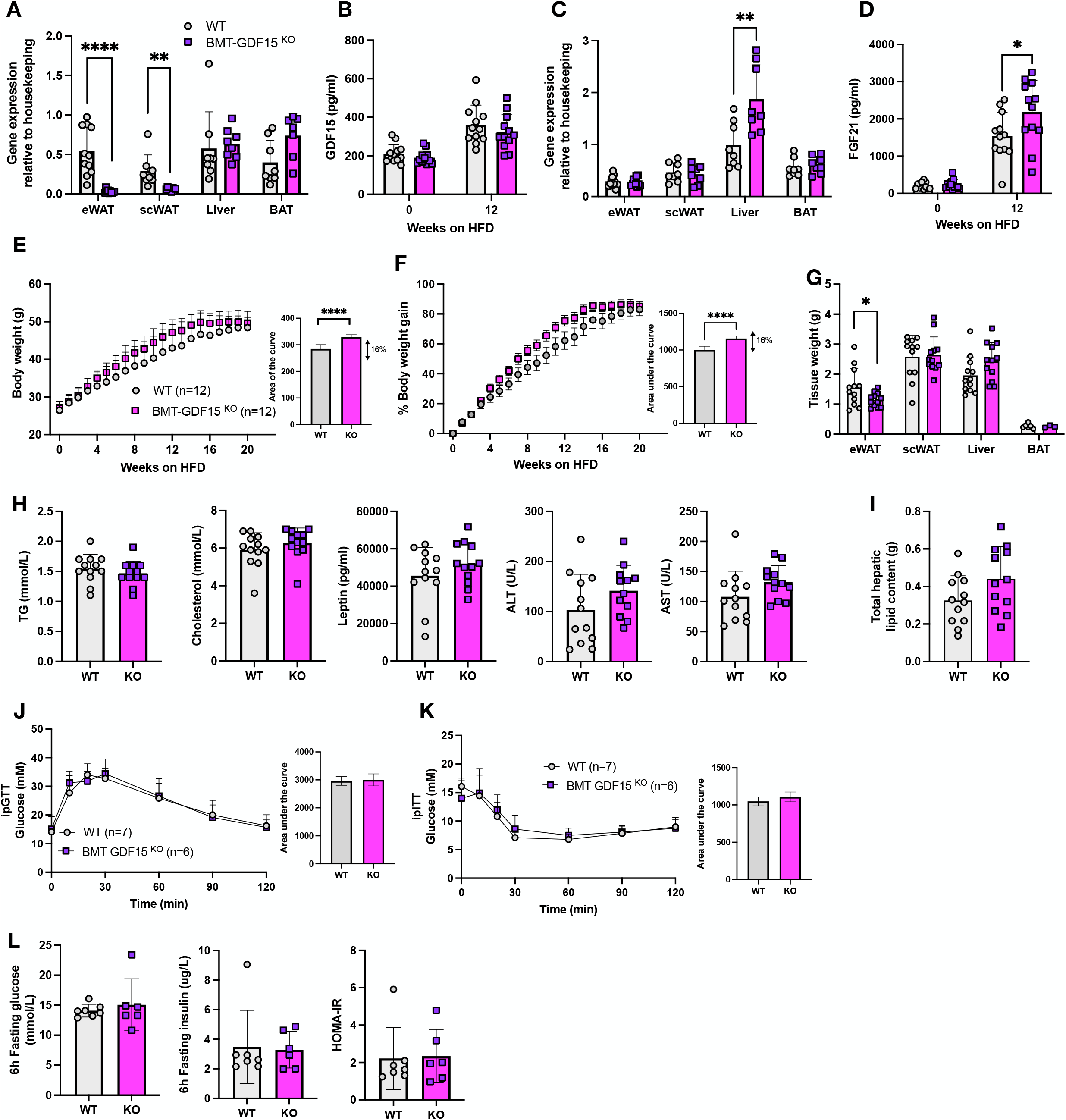
Phenotypic characterization of bone marrow deleted GDF15 knockout mouse on a high fat diet. (A) Gdf15 mRNA expression in tissues from 24 week old high fat diet (HFD) fed WT and BMT-GDF15^KO^ mice (n=8-12). (B) Plasma GDF15 levels from random fed mice at the onset (week 0) and after 12 weeks of HFD feeding (n=12). (C) Fgf21 mRNA expression from 24 week HFD fed WT and BMT-GDF15^KO^ mice (n=8-12). (D) Plasma FGF21 levels from random fed mice at the onset (week 0) and after 12 weeks of HFD feeding (n=12). (E and F) Body weight and percent body weight gain of WT and BMT-GDF15^KO^ mice fed a 60% HFD; Inset, area of curve analysis of body weight over time and area under the curve analysis of % body weight gain over time. (G) Weight of epididymal white adipose tissue (eWAT), subcutaneous white adipose tissue (scWAT), liver and brown adipose tissue (BAT) harvested at the end of the study, 24 weeks of HFD (n=3-12). (H) Plasma Triglyceride (TG), cholesterol, leptin, alanine transaminase (ALT) and aspartate transaminase (AST) from random fed mice, after 16 weeks of HFD feeding (n=12). (I) Weight of total lipids isolated per mg of liver, normalized to total liver weight (n=12). (J and K) Blood glucose levels during ip glucose tolerance test (GTT) and insulin tolerance test (ITT) after 16 weeks of HFD feeding. Inset, area under the curve analysis of glucose over time. (L) Blood glucose, plasma insulin and HOMA-IR levels from 6h fasted mice after 16 weeks of HFD feeding (n=7,6). All data are means ± S.D */**/***/**** - p<0.05/0.01/0.001/0.0001

Collectively, these data suggest that myeloid cells are the main producer of adipose tissue GDF15 but contribute very little to the rise in circulating GDF15 concentrations seen in HFD fed mice. An implication of this finding is that white adipose tissue is likely to be a minor contributor to the rise in circulating GDF15 concentrations associated with weight gain. In keeping with this observation, plasma GDF15 concentrations were only weakly associated with white adipose tissue *GDF15* mRNA in the human TwinsUK cohort referred to previously (β=0.08; *P*=0.02) **(Figure 4 supplemental)**.

### Characterisation of liver specific GDF15 null mice

In order to evaluate the contribution of hepatocyte derived GDF15, we used an Albumin driven Cre line (Alb-Cre) to generate hepatocyte-specific GDF15 null mice. Effective deletion of GDF15 was confirmed within Alb-GDF15^KO^ liver tissue where there was a 90% reduction in GDF15 mRNA expression **(Figure 5A)**. Furthermore, in this instance plasma GDF15 was significantly reduced (∼50%) in HFD fed mice **(Figure 5B)**, establishing the liver as a major source of circulating GDF15 in the HFD fed state. Alb-GDF15^KO^ mice also displayed elevated circulating FGF21 levels upon HFD feeding, but FGF21 mRNA expression in the liver was unchanged **(Figure 5C-D)**. Similarly to the macrophage deletors, Alb-GDF15^KO^ mice were slightly heavier on the HFD compared to the WT control groups **(Figure 5E-F)** though leptin concentrations were similar to those of WT littermates **(Figure 5H)**. However, the Alb-GDF15^KO^ mice did manifest larger livers **(Figure 5G)**, higher plasma lipids (TG, cholesterol) and liver transaminases (ALT, AST) **(Figure 5H)**. Histologically, Alb-GDF15^KO^ livers appeared to display similar steatosis compared to WT control mice **(Figure 5 supplemental)**, though total hepatic lipid content was significantly higher as the livers were heavier **(Figure 5I)**. Metabolically, Alb-GDF15^KO^ mice showed minor impairments in glucose and insulin tolerance as well as higher insulin concentrations **(Figure 5J-L)**.

**Figure 5:**
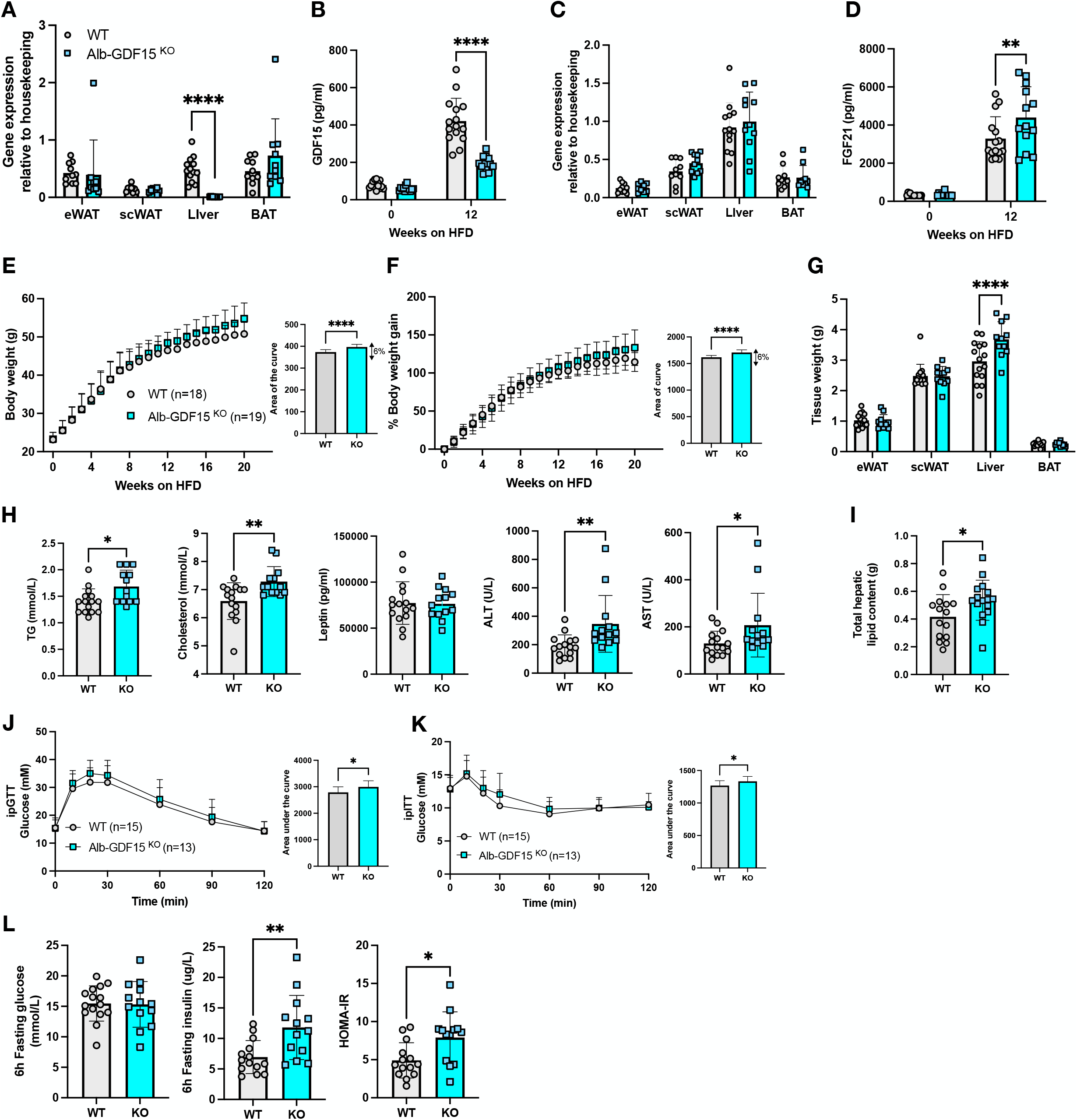
Phenotypic characterization of Alb-Cre mediated GDF15 hepatocyte knockout mouse on a high fat diet. (A) Gdf15 mRNA expression in tissues from 24 week old high-fat diet (HFD) fed wild type (WT) and Alb-GDF15^KO^ mice (n=10-13). (B) Plasma GDF15 levels from random fed mice at the onset (week 0) and after 12 weeks of HFD feeding (n=15,13). (C) Fgf21 mRNA expression from 24 week HFD fed WT and Alb-GDF15^KO^ mice (n=10-13). (D) Plasma FGF21 levels from random fed mice at the onset (week 0) and after 12 weeks of HFD feeding (n=15,13). (E and F) Body weight and percent body weight gain of WT and Alb-GDF15^KO^ mice fed a 60% HFD; Inset, area of curve analysis of body weight over time and area under the curve analysis of % body weight gain over time (G) Weight of epididymal white adipose tissue (eWAT), subcutaneous white adipose tissue (scWAT), liver and brown adipose tissue (BAT) harvested at the end of the study, 24 weeks of HFD (n=15,11). (H) Plasma triglyceride (TG), cholesterol, leptin, alanine transaminase (ALT) and aspartate transaminase (AST) from random fed mice, after 16 weeks of HFD feeding (n=15,13). (I) Weight of total lipids isolated per mg of liver, normalized to total liver weight (n=15,16) (J and K) Blood glucose levels during ip glucose tolerance test (GTT) and insulin tolerance test (ITT) after 16 weeks of HFD feeding. Inset, area under the curve analysis of glucose over time. (L) Blood glucose, plasma insulin and HOMA-IR levels from 6h fasted mice, after 16 weeks of HFD feeding (n=14,13). All data are means ± S.D */**/***/**** - p<0.05/0.01/0.001/0.0001

### Characterisation of GDF15 /FGF21 double KO mice

Similarly to GDF15, circulating FGF21 has also been shown to be largely derived from the liver in obese HFD fed mice (Markan et al., 2014) raising the question of why two stress responsive hormones are secreted in this context of overnutrition. Furthermore, we had noted increased FGF21 concentrations in the plasma of GDF15 null mice suggesting that this might be attenuating the impact of GDF15 deficiency on HFD associated weight gain. In order to test this hypothesis we next studied mice lacking both GDF15 and FGF21 i.e. ‘double knockouts’. The GDF15/FGF21 double knockout (DKO) strain displayed normal viability and fertility and had no gross abnormalities. Loss of both hormones in DKO mice was confirmed by checking mRNA expression in the liver **(Figure 6A-B)** and circulating concentrations **(Figure 6C-D)**. Interestingly, as seen for circulating FGF21, plasma GDF15 in FGF21KO mice was significantly higher than in wild-type littermates within 8 weeks of commencing a HFD diet and then further elevated after 16 weeks of HFD feeding **(Figure 6C)**. DKO mice were heavier than their WT littermates, but their weight was not significantly different from GDF15 KO mice **(Figure 6E)**. In contrast, in our facility, FGF21 KO mice were leaner than WT mice, as had been reported previously by Li et al. (Li et al., 2018) **(Figure 6E-F)**. Similarly to the GDF15 KOs, the DKO mice displayed impaired glucose and insulin tolerance (as reflected by AUC) **(Figure 6G-H)**. However, higher fasting glucose, insulin and thus HOMA IR scores suggested that they were significantly more insulin resistant than the GDF15 null mice **(Figure 6I)**. In contrast, FGF21 KO mice displayed relatively normal glucose tolerance and circulating insulin concentrations **(Figure 6I)**.

**Figure 6:**
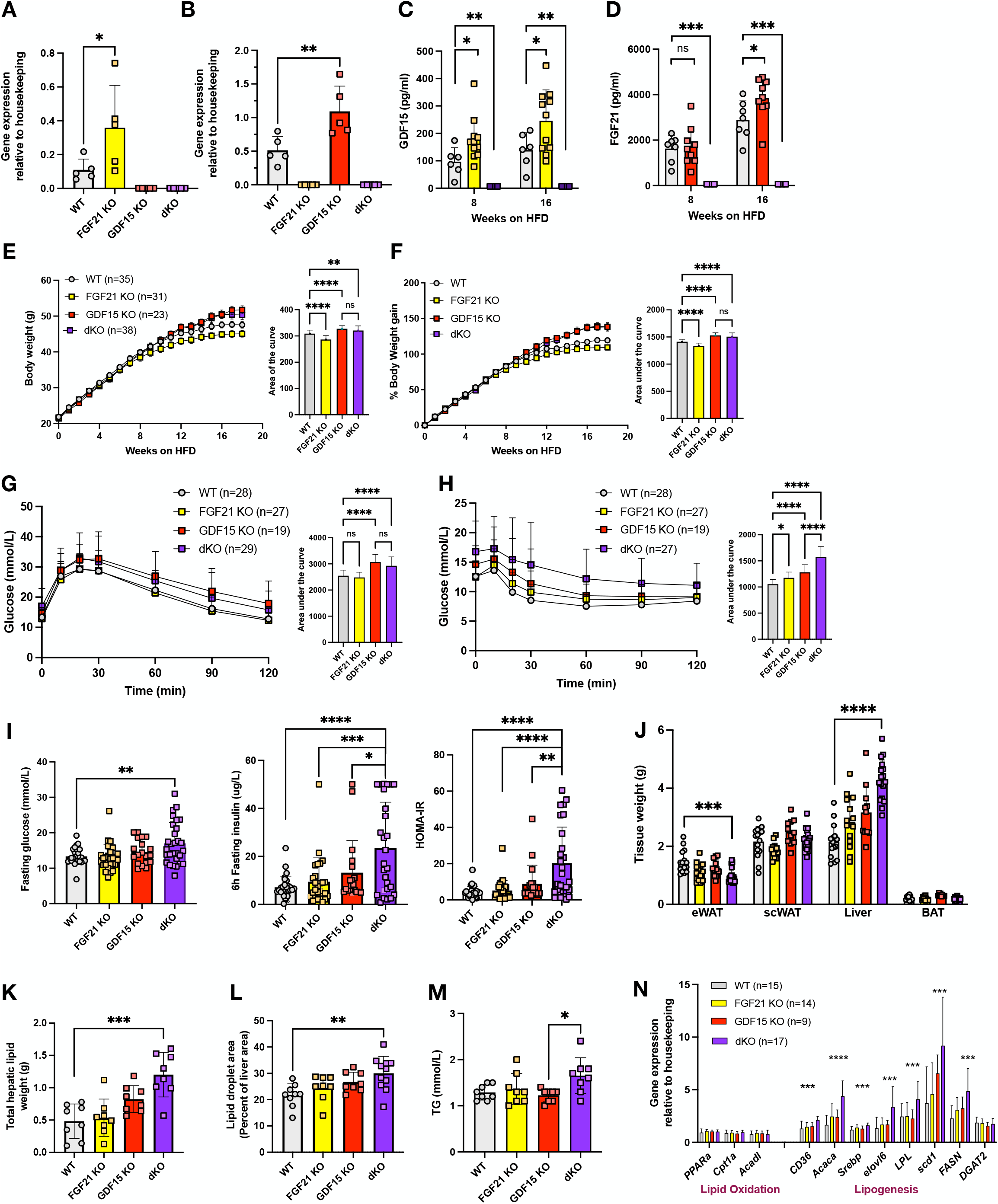
Phenotypic characterization of DKO knockout mice on a high fat diet. (A) Liver Gdf15 and (B) Fgf21 mRNA expression from 24 week old high fat diet (HFD) fed WT, FGF21 KO, GDF15 KO and FGF21/GDF15 double knockout (dKO) mice (n=5) (C and D) Plasma GDF15 and FGF21 from random fed mice at indicated time after high fat feeding (n=3-11). (E and F) Body weight and percent body weight gain of WT, FGF21 KO, GDF15 KO and dKO mice fed a 60% HFD; Inset, area of curve analysis of body weight over time and area under the curve analysis of % body weight gain over time (G and H) Blood glucose levels during ip glucose tolerance test (GTT) and insulin tolerance test (ITT) after 16 weeks of HFD feeding. Inset, area under the curve analysis of glucose over time. (I) Blood glucose, plasma insulin and HOMA-IR levels from 6h fasted mice after 16 weeks of HFD feeding (n=19-29). (J) Weight of epididymal white adipose tissue (eWAT), subcutaneous white adipose tissue (scWAT), liver and brown adipose tissue (BAT) harvested at the end of the study, 24 weeks of HFD (n=11-17). (K) Weight of total lipids isolated per mg of liver, normalized to total liver weight (n=8). (L) Lipid droplet area (Percent liver area) determined from histological analyses of H&E stained liver sections (n=8-11). (M) Plasma triglycerides (TG) from random fed mice, after 24 weeks of HFD feeding (n=8). (N) Hepatic mRNA expression of genes involved in lipid metabolism (n=9-17). All data are means ± S.D */**/***/**** - p<0.05/0.01/0.001/0.0001

Despite their body weight being similar to that of the single GDF15 KOs, the most prominent phenotype of the DKO mice was their increased liver size **(Figure 6J)**. This was associated with increased hepatic lipid content and elevated circulating TGs **(Figure 6K-M and Figure 6A supplemental).** Furthermore, plasma cholesterol, NEFA, AST and ALT levels were also significantly higher in the DKOs compared to the other genotypes **(Figure 6B supplemental)**. Gene expression analysis revealed increases in genes encoding proteins involved in *de novo* lipogenesis (DNL) whereas genes involved in fat oxidation were expressed at similar levels to the other genotypes studied (**Figure 6N)**. This is likely to relate to the higher insulin concentrations in the DKO mice.

## Discussion

GDF15 was identified more than 20yrs ago as a gene that was upregulated in ‘activated macrophages’ but has since been shown to be elevated in a range of human disease states (Breit et al., 2021; Keipert and Ost, 2021; Wang et al., 2021). Our original interest in understanding GDF15 biology in the context of weight gain and metabolic disease related to three key observations: i) GDF15 is elevated in human and rodent models of obesity (Carballo-Casla et al., 2021; Vila et al., 2011; Xiong et al., 2017); ii) transgenic overexpression or pharmacological treatment with GDF15, suppresses food intake and reduces body weight in mice (Chrysovergis et al., 2014; Johnen et al., 2007; Macia et al., 2012; Xiong et al., 2017); and iii) this effect is mediated exclusively via the hindbrain restricted expression of the GFRAL receptor (Emmerson et al., 2017; Mullican et al., 2017). Whilst significant progress has been made in understanding the mechanism by which GDF15 leads to suppression of food intake and modulation of body weight (Emmerson et al., 2017; Macia et al., 2012; Mullican et al., 2017), the cellular source of GDF15 in this context remains unclear.

Here we first sought to determine i) the source of circulating GDF15 in HFD fed mice and ii) if selective deletion of this source replicates the more obese phenotype observed in global GDF15 KO mice on a HFD. We previously showed that expression of GDF15 mRNA is increased in white and brown adipose tissue and in the liver in mice fed a HFD for up to 18 weeks, whereas its expression in skeletal muscle in this context is very low and does not differ from that of chow fed mice (Patel et al., 2019). We used RNA scope to reveal the cellular source of GDF15 in WAT to be almost exclusively macrophages and in the liver, to be the hepatocytes themselves. In brown fat, we see a weak signal in selected adipocytes as well as in stromal macrophages. In keeping with these murine observations, we observed a strong correlation between GDF15 mRNA and macrophage markers (CD68 and EMR1) in human white fat samples from two large independent cohorts. It is well established that the number of macrophages in white adipose tissue increases with weight gain so these findings could either simply relate to the higher number of macrophages present in the tissue and/or increased expression within individual macrophages.

Interestingly, despite this evidence suggesting that macrophages account for the increase in GDF15 mRNA in WAT associated with weight gain in humans and mice, our data does not suggest that macrophages contribute a great deal to circulating GDF15 concentrations in this context. This is borne out by the very modest impact of both LysM Cre driven GDF15 deletion in macrophages and the reconstitution of hematopoietic lineage cells from GDF15 null mice in irradiated wild type mice, on serum GDF15 levels in HFD fed mice. Furthermore, this and the hepatocyte specific deletion of GDF15, indicate that Kuppfer cells (liver resident macrophages) are not a major souce of GDF15. In humans, we also noted that plasma GDF15 levels correlate weakly with WAT GDF15 expression. Nevertheless, we did see a very modest tendency for the above mouse lines to gain more weight than wildtype controls when fed a HF diet.

When GDF15 was deleted in hepatocytes using the Alb-Cre promoter, we observed a more substantial impact on GDF15 concentrations in mice gaining weight on a HFD, suggesting that the liver is likely to be the major source of GDF15 in this context. These data are also consistent with a recent human study which showed that plasma GDF15 levels correlated strongly with NAFLD progression (Govaere et al., 2020). We have previously shown that activation of the integrated stress response is likely to be mediating the increase in GDF15 mRNA expression in the liver, highlighting the ‘stress’ imposed on hepatocytes by overfeeding. Hepatocytes appear to be a major site of ectopic lipid accumulation when the capacity of white adipocytes to store surplus energy is overloaded, and these data suggest that this results in a stress response in the hepatocytes, one of the consequences of which is increased GDF15 secretion. In contrast to this stress response in hepatocytes, skeletal muscle does not appear to manifest a similar response in high fat fed mice. We are also not aware of activation of the ISR having been demonstrated in skeletal muscle in obese humans.

In terms of the impact of the changes in circulating GDF15 concentrations on body weight, we saw very modest changes which did not necessarily correlate with the changes or lack thereof in GDF15 concentrations. This is broadly consistent with the rather subtle impact of global GDF15 deficiency on body weight. Interestingly we did observe higher FGF21 expression in the GDF15 null mice and in some of the tissue specific GDF15 knockout lines. In order to determine if this response might be mitigating the impact of GDF15 deficiency on body weight we proceeded to study mice deficient for both hormones. In this context, we also showed that circulating concentrations of GDF15 are higher in FGF21 deficient mice. Whilst body weight gain of the double KO (DKO) mice was greater than that of wildtype littermates in response to HFD feeding, the weight of the DKO mice was no higher than that of GDF15 null mice (Figure 6C). Nevertheless, the GDF15/FGF21 DKO mice did appear to be significantly more insulin resistant than wild-type mice and, both GDF15 and FGF21 ‘single knockouts’. Fatty liver disease was also significantly worse in the DKO mice and was accompanied by higher plasma triglycerides and liver transaminases.

In this study, we have identified the liver as a major source of plasma GDF15 upon high-fat feeding in mice. However, the data suggest that additional tissues probably also contribute small amounts towards circulating GDF15 concentrations in this paradigm. This study also highlighted the subtle impact of the GDF15/GFRAL axis on body weight regulation in HFD fed mice. The HFD-induced increase in plasma GDF15 is relatively modest in magnitude and gradual in its onset when compared with that associated with acute toxic stressors or some forms of cancer. The latter clearly reduce food intake and body weight, whereas the HFD induced changes in GDF15 appear to have a minor anorectic effect which is reflected in the discordant reports of body weight phenotypes observed in GDF15 and GFRAL KO mice (Emmerson et al., 2017; Hsu et al., 2017; Mullican et al., 2017; Tran et al., 2018; Yang et al., 2017). Interestingly, this study did reveal an additive impact of GDF15 and FGF21 deletion on liver steatosis and insulin resistance despite the double knockout mice having similar body weights to the GDF15 knockouts. Whilst lipid synthesis genes were upregulated this phenotype is also likely to reflect additional consequences of the loss of GDF15 and FGF21 signalling. Recent work by Katsumura et al (Katsumura et al., 2022) clearly showed that increasing FGF21 and GDF15 together reduces food intake and increases energy expenditure, in keeping with prior work on their respective mechanisms of action (Hsu et al., 2017; Owen et al., 2014).

Collectively the data indicate that similarly to the situation with FGF21, the liver is a major source of circulating GDF15 in the context of high fat feeding. The fact that Alb-Cre driven deletion of GDF15 does not entirely alleviate the HFD induced rise in GDF15 suggests that other tissues are also likely to be contributing smaller amounts. Importantly however, we have never seen an increase in GDF15 or FGF21 in skeletal muscle in the context of HF feeding despite other data suggesting that muscle can secrete both hormones in the face of mitochondrial stress for example (Chung et al., 2017; Forsström et al., 2019; Ost et al., 2020). GDF15 deficiency does appear to result in an increase in high fat feeding induced weight gain though the size of this effect is modest. We interpret this as reflecting the primary role of GDF15 as a stress responsive hormone which has evolved to reduce eating and promote rest in the context of more acute insults such as might occur after ingesting a toxin. FGF21 is also highly expressed in the liver in high fat fed mice and does appear to have an additive impact on alleviating fatty liver and preserving insulin sensitivity, but the net impact on body weight is no greater than that associated with GDF15 deficiency alone.

## Supporting information

Patel_Haider_supplemental

## Acknowledgements

The authors thank the Clinical Biochemistry Assay Lab, Disease Model Core (DMC), Histopathology Core and Imaging Core facilities at IMS-MRL, University of Cambridge, for experimental support for this study. This work was supported by the Medical Research Council (MRC) Metabolic Diseases Unit [MC_UU_00014/5] and the Wellcome Trust Major Award [208363/Z/17/Z]. J.A.T is supported by the MRC Metabolic Diseases Unit (MC_UU_00014/1) and by a NIHR Clinical Lectureship (CL-2019-14-504). D.B.S (WT 219417) and S.O. are supported by the Wellcome Trust (WT 214274/Z/18/Z), the MRC Metabolic Disease Unit (MC_UU_00014/1), and the NIHR Cambridge Biomedical Research Centre and NIHR Rare Disease Translational Research Collaboration. S.V. and G.B. are supported by The British Heart Foundation (RG/18/7/33636), and by the MRC (MC_UU_00014/2). M.B and E.G-J.was supported by the Deutsche Forschungsgemeinschaft (DFG, German Research Foundation) through CRC 1052, project number 209933838, subproject B1 to M.B.and by Deutsches Zentrum für Diabetesforschung (DZD, Grant: 82DZD00601) to M.B and E.G-J. KSS acknowledges funding from the Medical Research Council (MR/M004422/1 and MR/R023131/1). TwinsUK is funded by the Wellcome Trust, Medical Research Council, European Union, Chronic Disease Research Foundation (CDRF), Zoe Global Ltd and the National Institute for Health Research (NIHR)-funded BioResource, Clinical Research Facility and Biomedical Research Centre based at Guy’s and St Thomas’ NHS Foundation Trust in partnership with King’s College London.

## Competing interests

S.O. undertakes remunerated consultancy work for Pfizer, AstraZeneca, GSK. MB received fees for lectures and consultancy from Amgen, AstraZeneca, Bayer, Lilly, Novartis, Novo Nordisk and Sanofi.

## STAR METHODS

### Contact for reagent and resource sharing

Further information and requests for resources and reagents should be directed to and will be fulfilled by the Lead Contact, David Savage.

### Animal husbandry

Mice were maintained in ventilated cages with group housing (2-4 per cage), unless specified otherwise for indirect calorimetry and food intake experiments, on a 12 h light/12 h dark cycle (lights on 06:00–18:00), in a temperature-controlled (20-24 °C) facility, with *ad libitum* access to food and water. During the experimental protocol, all mice were fed either *ad libitum* or fasted as stated prior to some tests. All animal studies were performed on male mice and carried out at two facilities at the University of Cambridge, UK. This research was regulated under the Animals (Scientific Procedures) Act 1986 Amendment Regulations 2012 following ethical review by the University of Cambridge Animal Welfare and Ethical Review Body (AWERB).

### Mouse models

Mice carrying the GDF15 knockout-first “tm1a” allele [C57BL/6N-Gdf15^tm1a(KOMP)Wtsi/H^] were obtained through the IMPC, from the Harwell production centre (https://www.mousephenotype.org/data/alleles/MGI:1346047/tm1a%2528KOMP%2529Wtsi.). A “conditional ready GDF15 tm1c” allele [C57BL/6N-Gdf15^tm1c(KOMP)Wtsi^/H] expressing mouse was generated in house. Briefly, one-cell stage embryos (obtained from super-ovulated wild type C57Bl/6N females fertilised *in vitro* with sperm from homozygous Gdf15 Tm1a male) were injected into the pronucleus with 100ng/ul StemMACS Flp Recombinase mRNA (Miltenyi Biotec) then transferred into the uteri of pseudo pregnant recipient females (F1 hybrids from C57Bl/6J female x CBA/Ca male crosses). Mice from the transfer (F1 mice) were analysed for the presence of the Gdf15 Tm1c and Tm1a alleles. The F1 founders were crossed twice with wild type C657Bl/6N mice before establishing the Gdf15 Tm1c and Gdf15 Tm1a colonies. The GDF15 Tm1c mouse model contains loxP sites flanking exon 2 of GDF15 gene (see https://www.mousephenotype.org/data/alleles/MGI:1346047/tm1c(KOMP)Wtsi). FGF21^-/-^ mice (B6N; 129S5-Fgf21tm1Lex/Mmucd) were generated on a C57BL/6N background using sperm obtained from MMRRC (https://www.mmrrc.org/catalog/sds.php?mmrrc_id=32306). Gdf15/Fgf21 DKO mice and wild-type littermates were obtained from het x het breeding set-ups with all strains maintained on a C57BL6/N background. Genotyping to confirm derivation of all mouse lines was done by PCR using the primers described in the KRT.

Myeloid-specific deletions of GDF15 were generated using two separate strategies:

1. GDF15^fl/fl^ (GDF15 Tm1c) mice were bred to transgenic *Lyz2*^Cre/+^ mice carrying Cre recombinase under the control of the LysM promoter (kindly provided by T.Vidal-Puig, Institute of Metabolic Sciences, Univ of Cambridge). This line was maintained on a mixed C57BL6N/J background. Genotypes were confirmed by PCR for the presence of Cre and for the detection of the floxed GDF15 allele. GDF15^fl/fl^ and GDF15^fl/fl^ LysM-Cre were used in this study.
2. 24 x C57BL/6N wild type mice (4-6wks of age) were purchased from Charles River, Italy and were allowed to acclimatize for one week before being irradiated. Mice received two separate doses of 5.5 Gy of radiation using Caesium 60 source with a 4 h gap in between doses. 1-2 hours post irradiation, donor bone marrow cells (10 million/mouse) from either male GDF15 KO Tm1a or wild type littermates were injected into the tail veins of the irradiated mice. The mice were then housed under standard conditions for one month, monitored and weighed regularly until 12 weeks of age prior to experimentation.

Hepatocyte-specific GDF15 KO mouse were generated by breeding GDF15^fl/fl^ (GDF15 Tm1c) to *Alb*^Cre/+^ mice carrying Cre recombinase under the control of the albumin promoter (kindly provided by A. Kaser, Univ of Cambridge). This line was maintained on a C57BL/6N background and genotypes were confirmed by PCR for the presence of Cre as well as floxed GDF15 allele. GDF15^fl/fl^ and GDF15^fl/fl^ Alb-cre were used in this study.

### Mouse studies

Starting at the age of 5–6-weeks, mice were fed either a control chow (R105-25, Safe Diets) or a 60 % high fat diet (D12492i, Research Diets) for a period of 14-26 weeks. For all cohorts, the mice were weighed weekly and body composition was determined every 4 weeks by Time-Domain Nuclear Magnetic Resonance (TD-NMR) using a Minispec Live Mice Analyzer (LF50, Bruker). Tail blood samples were collected into heparinized micro blood tubes (01605-00, Hawksley), centrifuged at 13,000 x g for 4 min and plasma was collected for the analysis of hormones or lipids as indicated in figures. The mice were euthanized at the end of the experiment with tissues harvested, weighed and frozen at -80 °C or fixed in 10% neutral buffered formalin (NBF).

### Glucose and insulin tolerance test

Glucose and insulin tolerance tests (GTT and ITT) were conducted on mice that were fasted for 6 h (08:00 -14:00) in clean cages. Mice were injected intraperitoneally with 1 g/kg glucose (for GTT) or 1.5 U/kg insulin (for ITT) and blood glucose (tail vein) was measured at the indicated times using a AlphaTRAK 2 meter. Blood samples (∼30 ul) were collected during a GTT for insulin measurements. Homeostatic model assessment for insulin resistance (HOMA-IR) was calculated as 6 h fasting glucose (mmol/l) x 6 h fasting insulin (ng/ml)/22.5.

### Serum, plasma and media analysis

Tail blood samples from mice were collected for plasma analysis. Mouse leptin and insulin were measured using a single-plex MesoScale Discovery assay kit (Rockville, MD, USA product code K152BYC-2 for leptin and K152BZC-3 for insulin). The assay was performed according to the manufacturer’s instructions using calibrators provided by MSD. Mouse GDF15 was measured using a DuoSet ELISA (R&D Systems product code DY6385) which had been modified to run as an electrochemiluminescence assay on the MesoScale Discovery assay platform. Mouse FGF21 was analysed using a Quantikine ELISA kit (R&D Systems, product code MF2100) following the manufacturer’s instructions. ALT, AST, Cholesterol and triglycerides were measured on the Siemens Dimension EXL autoanalyser using Siemens reagents and calibrators. Human GDF15 was also measured using a DuoSet ELISA (R&D Systems product code DY957) that had been converted to MSD format. Comprehensive Quality Control procedures were followed for all measurements and no results were reported without passing QC checks.

Mouse sample measurements were performed by the Cambridge MRC MDU Mouse Biochemistry Laboratory and human assays were provided by the NIHR Cambridge BRC Core Biochemical Assay laboratory (CBAL).

### Histology, immunohistochemistry and tissue architecture analysis

Tissues were dissected and placed into 10% formalin for 48 h at room temperature, transferred to 70% ethanol and embedded into paraffin. Five-micrometre sections were cut using a Leica microtome, mounted onto Superfrost Plus slides (Thermo Fisher Scientific) and stained for hematoxylin and eosin. Detection of mouse *Gdf15* and *Fgf21* mRNA was performed on formalin-fixed paraffin-embedded sections, obtained from 45% high-fat diet fed, wild-type C57BL6/J mice (Patel et al., 2019), using Advanced Cell Diagnostics (ACD) RNAscope 2.5 LS Reagent Kit-RED (no. 322150) and RNAscope LS 2.5 Probe Mm-Gdf15-O1 (no. 442948) (ACD) and and RNAscope LS 2.5 Probe Mm-Fgf21-C2 (no. 460938-C2). Slides were processed as previously described (Adriaenssens et al., 2019). Positive and negative controls were run in parallel each time

All slides were imaged using the Zeiss AxioScan.Z1 Slide Scanner at x20 (standard) or x40 magnification (RNAscope). For the RNAscope slides, three Z-stacks spanning a total of 2 μm were acquired and merged into a single extended depth of focus (EDF) image with maximum projection processing, and then sharpened using Unsharp Masking (ZEN Blue, Zeiss). Hepatic lipid droplet area was determined using the vacuole detection algorithm of the Halo software (Indica labs). Hepatic lipid area was expressed as a percentage of total hepatic tissue area scanned per section.

### Hepatic Lipid content measurement

Accumulation of hepatic lipid was determined using a modified Folch method. In brief, 25mg of frozen liver was homogenized in with 1.2 ml of a 2:1 ratio choloroform:methanol mixture (Sigma-Aldrich, Dorset, UK). Deionized water (240 μl) was then added to the mix, vortexed thoroughly, and samples centrifuged for 10 min at 16,100 *g* to generate a distinct organic and aqueous phase. The lower organic phase (500 μl) was collected into a preweighed glass tube and dried under a nitrogen stream with the lipid content weighed and normalised to total liver weight.

### Isolation of primary adipose stromal fractions

Epididymal adipose tissues were removed at the time of sacrifice and minced finely into small pieces and resuspended in 5ml Hanks’ Balanced Salt Solution (HBSS, H9269, Sigma), 0.1 g bovine serum albumin (BSA, A8806, Sigma), and 10 mg collagenase type II (C6885, Sigma). The tissue was completely disaggregated by incubation in a 37 °C shaker for approximately 15 min. The digested material was topped with 5 ml of ice-cold MACS buffer (2 mM EDTA, 0.5 % BSA in PBS) and allowed to settle for 5 min at room temperature. The adipocyte fraction floating on top was collected by pipetting, washed twice with MACS buffer and then snap frozen in Buffer RLT (Qiagen) for RNA extraction. The remainder of the digested solution (approx 9 ml) was filtered through a 100 µm nylon mesh cell strainer (Falcon 352360) and centrifuged at 400 x g for 5 min. The pellet containing the stromo-vascular fraction (SVF) was collected and washed once with MACS buffer and then resuspended with CD11b microbeads (130-049-601-Millteyni Biotec) in MACS buffer. The CD11b positive and negative cells (SVF) were separated and collected using MACS LS columns (130-042-401, Miltenyi Biotec) that were placed onto a magnetic field of a MACS separator (Miltenyi Biotec). The cells were centrifuged at 400 x g for 5 min and snap frozen in Buffer RLT (Qiagen) for RNA extraction.

### Culture and differentiation of bone-marrow derived macrophages

Femur and tibia bones from mice were isolated, cleaned and flushed with 10 ml of RPMI-1640 media (Sigma) through each bone using a 25G syringe. The flushed bone marrow cells were passed into a 100-μm cell strainer and centrifuged at 400 x g for 5 min and resuspended in macrophage differentiation medium [RPMI-1640 with 20% L929 conditioned medium, 10% heat-inactivated FBS and 100 U/ml penicillin-streptomycin (Thermo Fisher Scientific)]. Total bone-marrow cells were counted using a Countess II automated cell counter (Thermo Fisher) and seeded onto 10 cm non-culture treated plates (Falcon) at a density of 5×10^6^ cells per plate and maintained for 7 days maintained at 37 °C in a humidified atmosphere of 5 % CO2. On day 5 of differentiation, medium was removed, and 10 ml of fresh macrophage differentiation medium was added to each plate. On day 7 of differentiation, macrophages were detached using ice-cold PBS containing 1 mM EDTA and centrifuged at 400 g for 5 min. The cells were resuspended in macrophage differentiation medium and seeded onto 24 well plate for 24h prior to experiments.

To make L929 conditioned medium, L929 cells (CCL-1, ATCC) were seeded in DMEM supplemented with 10% heat-inactivated FBS, 100 U/ml penicillin-streptomycin and 2 mM L-glutamine (Sigma) at a density of 250,000 cells per 50 ml of medium per T175 tissue culture flask. Medium was harvested after 1 week of culture, and then 50 mL of fresh DMEM supplemented with 10% heat-inactivated FBS, 100 U/ml penicillin-streptomycin and 2 mM L-glutamine was added onto cells and harvested 1 week later. Batches obtained after the first and second weeks of culture were mixed at a 1:1 ratio, aliquoted and stored at -20 °C.

### Gene expression data for mouse studies

Following treatments, cells were lysed with Buffer RLT (Qiagen) containing 1% 2-Mercaptoethanol and processed through a Qiashredder with total RNA extracted using the RNeasy isolation kit according to manufacturer’s instructions (Qiagen). Meanwhile for mice, tissues were harvested and immediately snap frozen in liquid nitrogen and stored at -80 °C until further analysis. For RNA isolation, approximately 30-50 mg of tissue was placed in Lysing Matrix D tubes and homogenized in 800 µl TRI reagent (Sigma-Aldrich) using the Fastprep-24 Homogenizer for 30 sec at 4-6 m/s (MP Biomedical). The resultant supernatant was transferred to an RNAse free tube and 200 µl chloroform (Sigma) added. The samples were vortexed and centrifuged at 13,000 rpm for 15 min at 4 °C. The upper phase was then transferred to a RNAse free tube and mixed with equal volume of 70 % ethanol before loading onto RNA isolation spin columns (Qiagen). RNA was then extracted using the RNeasy isolation kit following the manufactureŕs instructions.

RNA concentration and quality was determined by Nanodrop. 400 - 600 ng of total RNA was treated with DNAase1 (Promega) and then converted to cDNA using Lunascript (NEB). Quantitative RT-PCR was carried out with either TaqMan™ Universal PCR Master Mix or SYBR Green PCR master mix on the QuantStudio 7 Flex Real time PCR system (Applied Biosystems). All reactions were carried out in either duplicate or triplicate and Ct values were obtained. Relative differences in the gene expression were normalized to expression levels of housekeeping genes B2M, 36b4 or RPL13a (geometrical mean) using the standard curve method. Primer sequences are shown in the key resources table.

### Human Studies

For the Twins cohort: Details of this study is described in (Buil et al., 2015). Briefly, the study included 856 female individuals of European ancestry recruited from the TwinsUK Adult twin registry. The study was approved by The St. Thomas’ Research Ethics Committee (REC). Volunteers gave informed consent and signed an approved consent form before the biopsy procedure.

For the obese cohort, 525 metabolically well-characterized participants of the Leipzig Obesity BioBank were recruited at four bariatric surgery centers in Leipzig, Karlsruhe, Dresden and Gera (all in Germany) (Table 1). All subjects underwent clinical phenotyping as described previously (Klöting et al., 2010; Langhardt et al., 2018; Rolle-Kampczyk et al., 2020). All subjects had a stable weight, defined as no fluctuations of > 2% of body weight for at least 3 months before surgery. According to American Diabetes Association (ADA) criteria (2014) 205 study participants (∼39%) were diagnosed with T2D. We defined the following exclusion criteria: i) thyroid dysfunction, ii) alcohol or drug abuse, iii) pregnancy and iv) treatment with thiazolidinediones. The study was approved by the ethics committee of the University of Leipzig (Approval numbers: 159-12-21052012 and 017-12-23012012). The study designs followed the Declaration of Helsinki and all participants gave written informed consent prior to participation.

**Table 1.** Anthropometric and metabolic characterization of the cohort.

|  | **BMI < 30 kg/m<sup>2</sup>**<br>**(n = 16)** | **BMI 30–40 kg/m<sup>2</sup>**<br>**(n = 51 )** | **BMI > 40 kg/m<sup>2</sup>**<br>**(n = 458)** |
| --- | --- | --- | --- |
| Age (years) | 65.09 ± 13.06 | 48.31 ± 11.66 | 47.02 ± 11.86 |
| Men/Women (n) | 12/4 | 12/39 | 119/339 |
| T2D (n) | 3 | 20 | 182 |
| Body weight (kg) | 78.88 ± 10.84 | 109.23 ± 12.67 | 143.29 ± 26.78 |
| Height (m) | 1.76 ± 0.09 | 1.70 ± 0.07 | 1.69 ± 0.09 |
| BMI (kg/m<sup>2</sup>) | 25.43 ± 2.31 | 37.51 ± 2.57 | 50.01 ± 7.30 |
| Body fat (%) | 22.88 ± 2.88 | 44.05 ± 9.76 | 49.59 ± 9.84 |
| Waist circumference (cm) | 95.17 ± 11.82 | 122.50 ± 9.19 | 145.56 ± 160.3 |
| Hip circumference (cm) | 98.33 ± 8.37 | 124.00 ± 11.31 | 148.06 ± 14.03 |
| WHR | 0.97 ± 0.09 | 0.99 ± 0.17 | 0.99 ± 0.10 |
| FPG (mmol/l) | 5.73 ± 0.62 | 5.98 ± 1.83 | 6.47 ± 2.45 |
| FPI (pmol/l) | 22.17 ± 11.72 | 111.86 ± 86.35 | 148.30 ± 110.35 |
| HbA1c (%) | 5.72 ± 0.48 | 6.05 ± 1.33 | 6.08 ± 1.16 |
| HOMA-Index | 0.77 ± 0.47 | 4.85 ± 4.24 | 6.07 ± 6.39 |
| Total Cholesterol (mmol/l) | 5.01 ± 0.76 | 5.20 ± 1.81 | 4.85 ± 1.09 |
| HDL-Cholesterol (mmol/l) | 1.21 ± 0.29 | 1.25 ± 0.30 | 1.17 ± 0.66 |
| LDL-Cholesterol (mmol/l) | 2.97 ± 0.37 | 3.33 ± 1.03 | 3.07 ± 0.95 |
| Triglycerides (mmol/l) | 1.23 ± 0.50 | 1.86 ± 1.21 | 2.12 ± 2.42 |
| CrP (mg/L) | 8.76 ± 12.60 | 7.70 ± 15.54 | 13.16 ± 18.03 |

Data are given as means ± SD. AT, adipose tissue; BMI, body mass index; CrP, C-reactive protein; FPG, fasting plasma glucose; FPI, fasting plasma insulin; HDL-C, high density lipoprotein cholesterol; LDL-C, low density lipoprotein cholesterol; SAT, subcutaneous adipose tissue; VAT, visceral adipose tissue; WHR, waist to hip ratio.

### Gene expression data for human studies

In the TwinsUK cohort, gene expression levels in subcutaneous adipose tissue were measured by RNA-Seq in 765 female twins from the TwinsUK cohort, as previously described (Buil et al., 2015). Briefly, punch biopsies were collected from a sun-protected area of the abdomen from each participant, from which adipose tissue was separated and RNA extracted. Gene expression was measured by RNA-Seq, with RNA-Seq reads aligned to the hg19 reference genome using STAR (Dobin et al., 2013) version 2.4.0.1, as fully described elsewhere (Glastonbury et al., 2019). Gene level counts were generated using the quan function of QTLtools (Delaneau et al., 2017) and Gencode version 19 (Frankish et al., 2019), and rank-based inverse normal transformation then applied to gene counts per million (CPMs) prior to all downstream analyses. Gene expression levels were then adjusted for RNA-Seq technical covariates using linear mixed effects models, with expression levels of each gene in turn treated as a dependent variable, with median insert size and mean GC content included as fixed effects, and date of RNA sequencing and RNA-Seq primer index as random effects.

In the obese cohort, paired samples of abdominal omental AT (visceral, VAT) and subcutaneous AT (SAT) were obtained from 525 Caucasian men (n = 143) and women (n = 382), who underwent open abdominal surgery as described previously (Langhardt et al., 2018; Rolle-Kampczyk et al., 2020). The age ranged from 18 to 80 years and body mass index (BMI) from 19 to 60 kg/m^2^. AT was immediately frozen in liquid nitrogen and stored at −80 °C. RNA was extracted from AT by using the RNeasy Lipid Tissue Mini Kit (Qiagen, Hilden, Germany), and qPCR was performed as described elsewhere (Mardinoglu et al., 2015; Stephane et al., 2006). Real-time quantitative PCR was performed with the TaqMan Assay predesigned by Applied Biosystems (Foster City, CA, USA) for the detection of human *GDF15* (Hs00171132_m1), *CD68* (Hs02836816_g1) and *GAPDH* (Hs 02786624_g1) mRNA expression in AT. All reactions were carried out in 96-well plates using the QuantStudio (TM) 6 Flex System Fast Real-Time PCR system. *GDF15* mRNA expression was calculated relative to *GAPDH* mRNA expression

### Estimation of adipose tissue cell type proportions

Adipose tissue cell type proportions were estimated from RNA-Seq data in TwinsUK using CIBERSORT (Newman et al., 2015), as reported previously elsewhere (Glastonbury et al., 2019). Estimated cell types included in our analysis were adipocytes, microvascular endothelial cells (MVEC) and macrophages.

## QUANTIFICATION AND STATISTICAL ANALYSIS

Cell and mouse quantitative data are reported as mean ± standard deviation (SD). As indicated in the figure legends, differences between means were assessed by two-tailed Student’s *t* tests or One-way ANOVA or Two-way ANOVA with sidak multiple comparisons test using either GraphPad Prism software (GraphPad, San Diego) or with SAS version 9.4, Cary, N. Carolina. Statistical significance was defined as *p* < 0.05.

Association analyses in TwinsUK cohort: Association of adipose tissue *GDF15* expression levels was assessed to *GDF15* serum levels, adipose tissue expression levels of macrophage markers *CD68* and *EMR1*, and adipose tissue estimated macrophage proportions, using linear mixed effects models. Participants with type 2 diabetes (n=32) were excluded from all analyses. Linear mixed effects models were fitted using the lmer function from the lme4 package (Bates et al., 2015) in R (R Development Core Team, 2013) version 3.5.1.

Rank-based inverse normal transformation was applied to serum GDF15 serum levels. GDF15 serum levels were treated as a dependent variable, with adipose tissue *GDF15* gene expression residuals, adjusted for technical covariates as previously described, as a predictor variable. Covariates included BMI and age as fixed effects, and family and zygosity as random effects. Family and zygosity are both random effects that permit identification of the family a twin belongs to, and their clonality (MZ/DZ status).

To assess association of adipose tissue *GDF15* expression levels with gene expression levels of macrophage markers *CD68* and *EMR1*, and adipose tissue estimated macrophage proportion, GDF15 gene expression residuals were treated as a dependent variable, and each of adipose tissue expression residuals of *CD68*, and *EMR1*, and adipose tissue estimated macrophage proportion, were used in turn as predictor variables. Models included the same fixed and random effect covariates as described for the GDF15 serum level model. For each analysis, the full model was then compared to a null model where the trait of interest was omitted, using a 1 degree of freedom ANOVA.

For obese human subject data, prior to statistical analysis, non-normally distributed parameters were logarithmically (ln) transformed to approximate a normal distribution. Results are expressed as mean ± SD. Linear regression analysis were used to assess the relationships between *GDF15* mRNA expression and phenotypic traits. Pearsońs correlation analyses were conducted using two-way bivariate correlations. Differences in *GDF15* mRNA expression between visceral and subcutaneous AT were assessed using the paired Student’s t-test or one-way ANOVA. Statistical analyses were performed using SPSS/PC+ for Windows statistical package (Version 27.0; SPSS, Chicago, IL, USA).

## Key Resources table

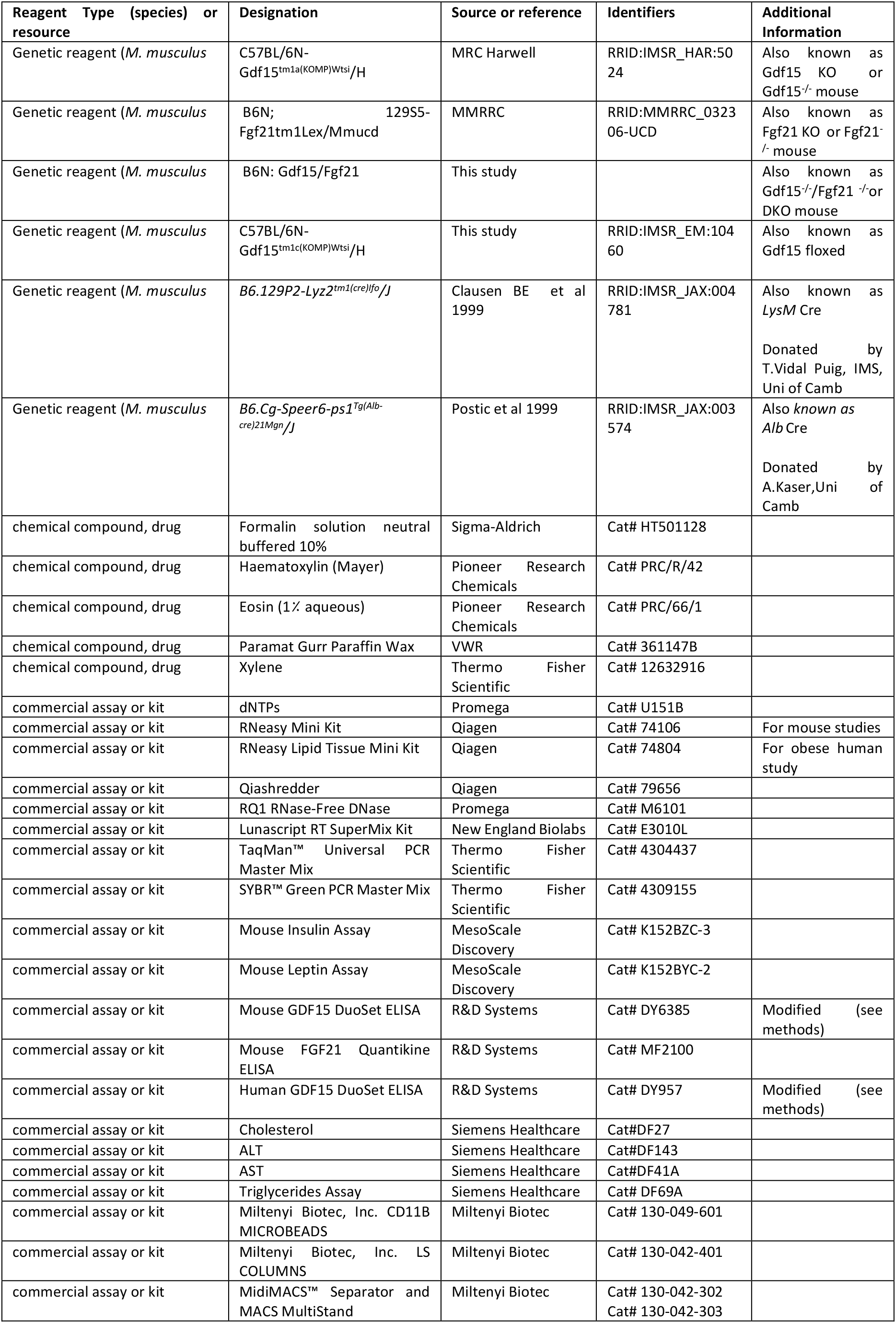
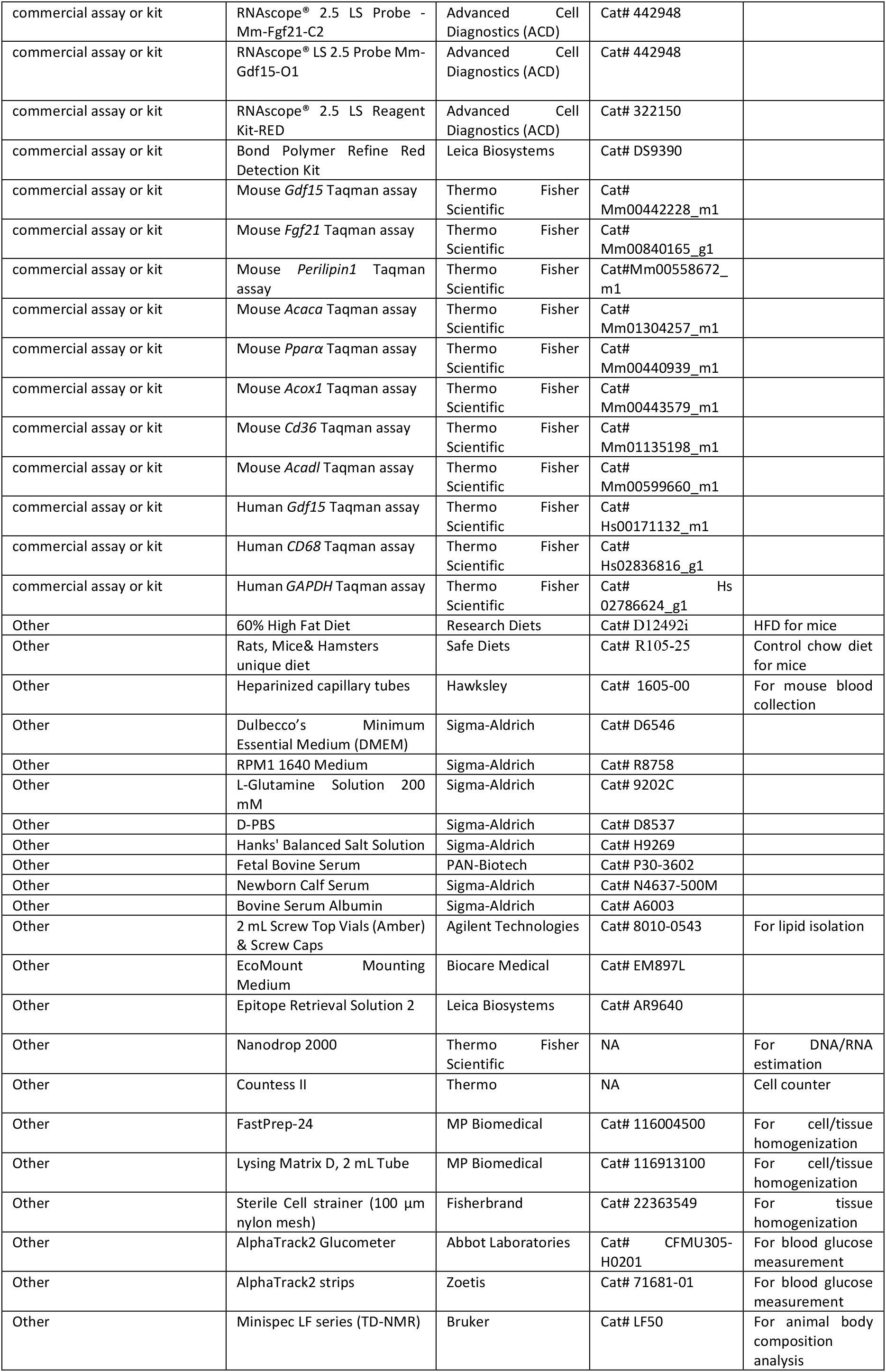
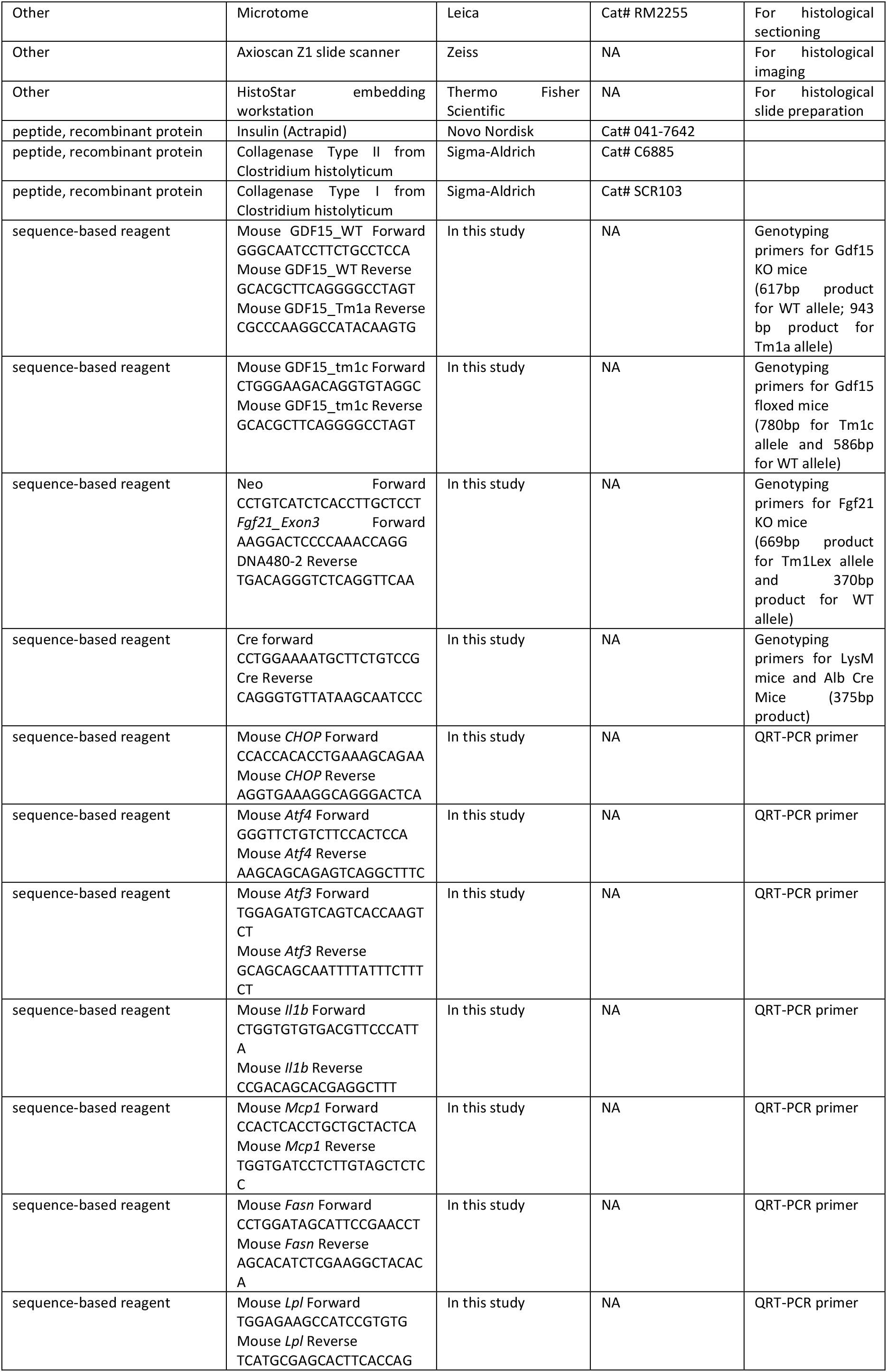
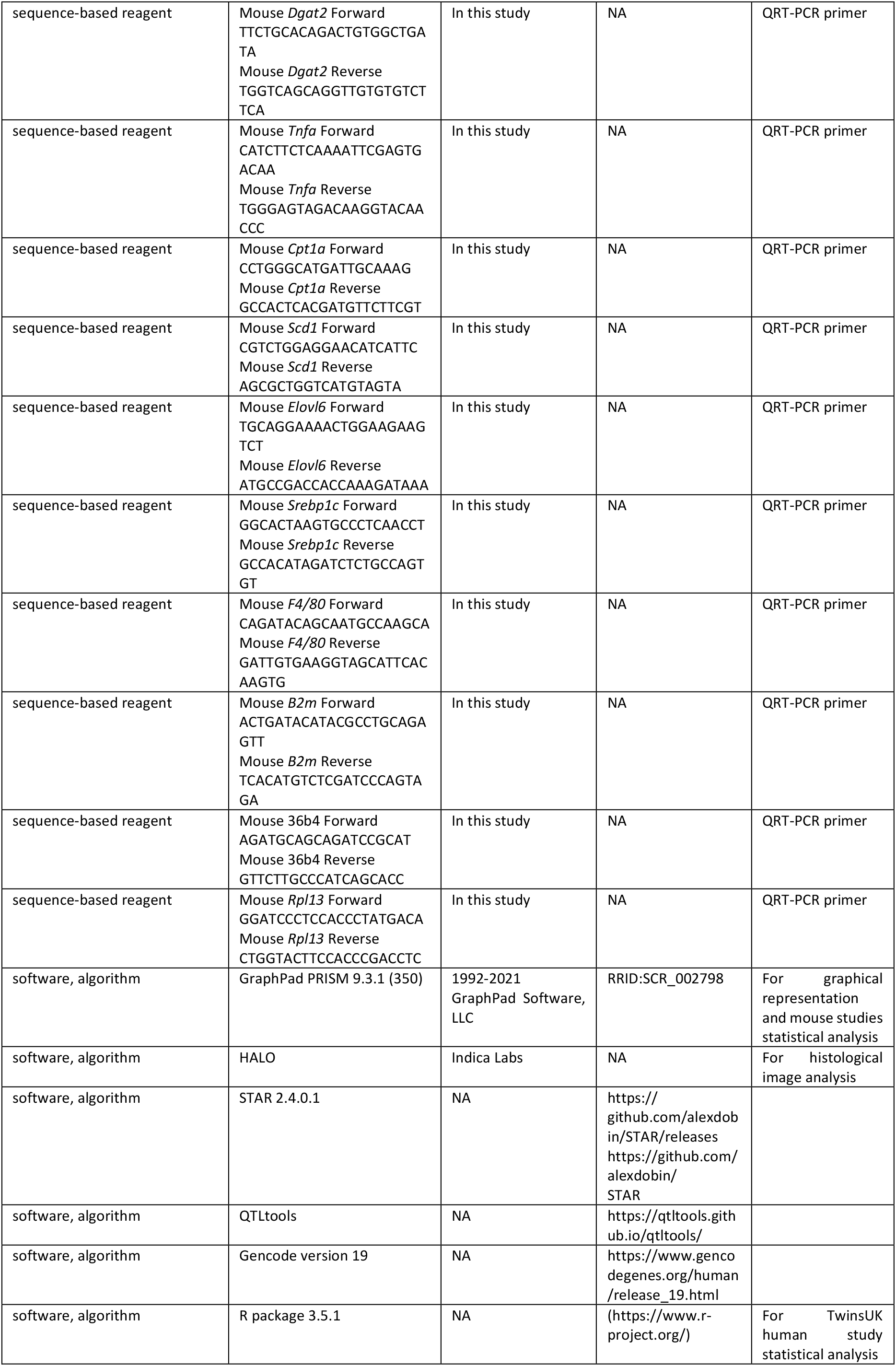
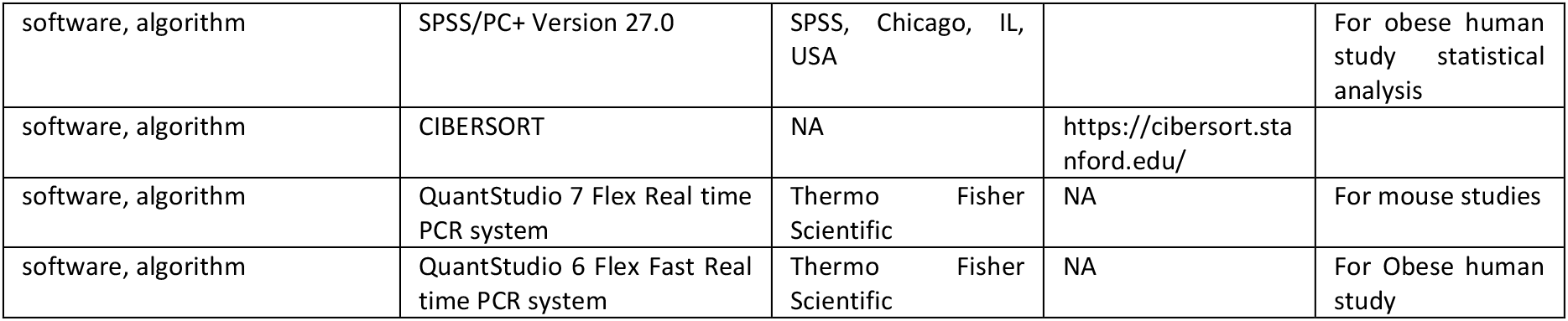

