## Supplementary material for "Endogenous GDF15 and FGF21 additively alleviate hepatic steatosis and insulin resistance in obese mice": Patel_Haider_supplemental

Cambridge, CB2 0QQ, UK.

[2] MRC Metabolic Diseases Unit, Wellcome-MRC Institute of Metabolic Science, University of Cambridge, Cambridge,UK.

[3] Department of Medicine, Division of Cardiovascular Medicine, University of Cambridge, Cambridge, United Kingdom;

[4] Medical Department III – Endocrinology, Nephrology, Rheumatology, University of Leipzig Medical Center, 04103 Leipzig, Germany

[5] Helmholtz Institute for Metabolic, Obesity and Vascular Research (HI-MAG) of the Helmholtz Zentrum München at the University of Leipzig and University Hospital Leipzig, Leipzig, Germany.

[6] Department of Twin Research and Genetic Epidemiology, King's College London, St Thomas' Campus, London, SE1 7EH, UK.

[7] East Midlands and East of England Genomic Laboratory Hub & Department of Histopathology, Cambridge University Hospitals NHS Foundation Trust, Cambridge, UK.

[8] These authors contributed equally

[9] Lead Contact

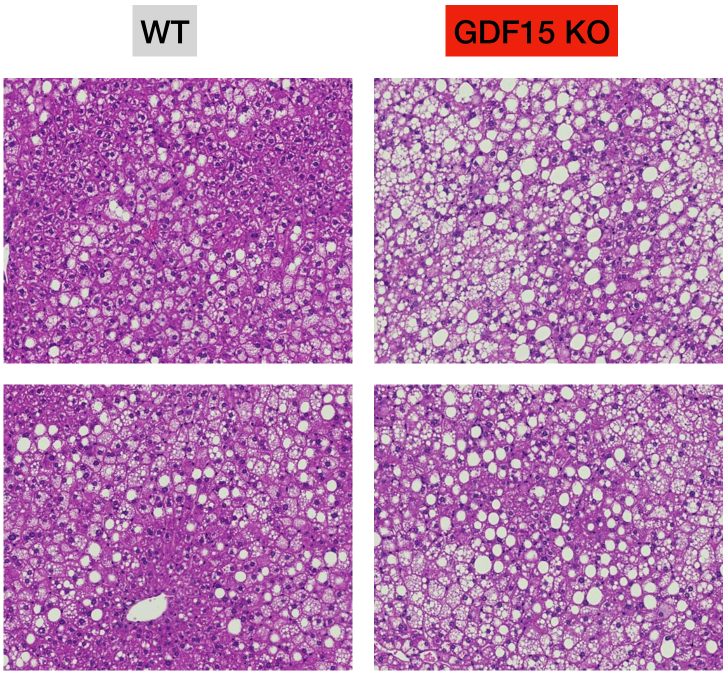


**Figure 1 Supplemental**
Representative images of haematoxylin/eosin stained liver sections from WT and GDF15KO mice after 25 weeks HFD feeding.

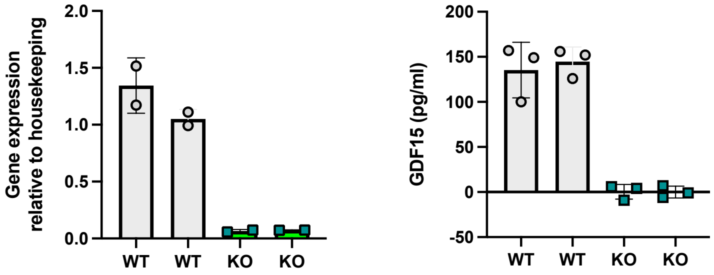


**Figure 3 Supplemental**

Gdf15 mRNA expression and secretion from WT and LysM-GDF15$^{KO}$ bone-marrow derived macrophages (BMDM).

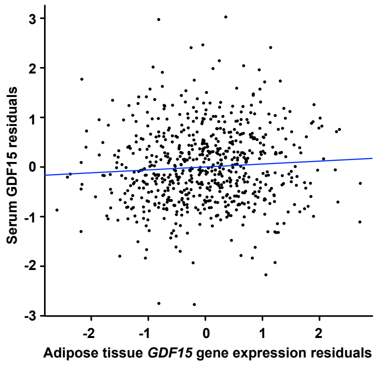


**Figure 4 Supplemental**

Human serum GDF15 levels show limited association with GDF15 gene expression levels in adipose tissue. Each point represents data from an individual. GDF15 serum levels were adjusted for age and BMI; plotted GDF15 gene expression residuals were adjusted for age, BMI and RNA-Seq technical covariates

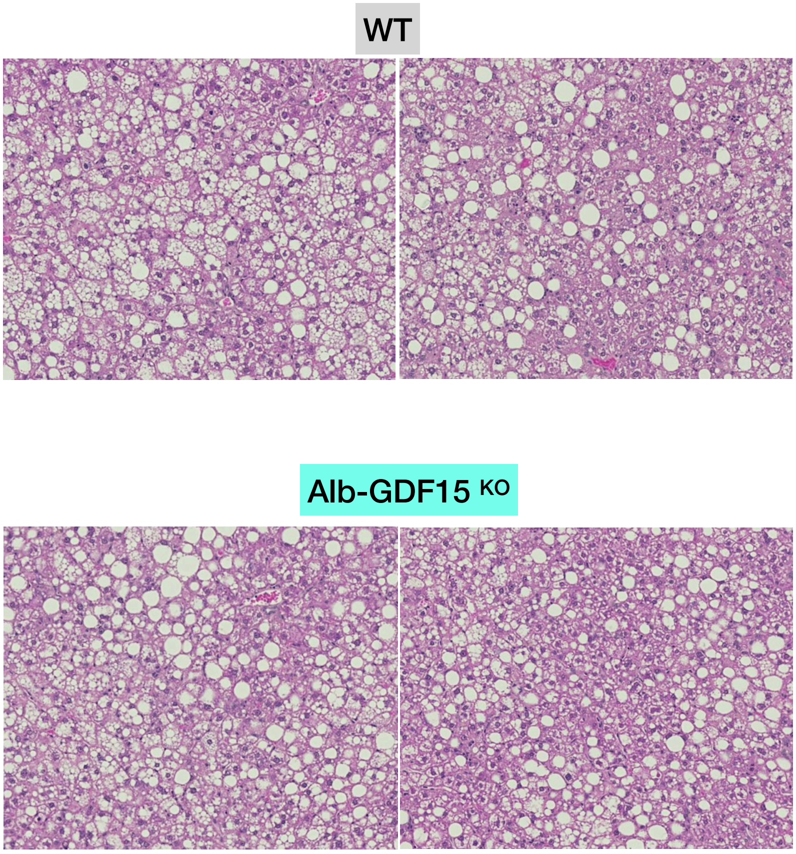


**Figure 5 Supplemental**

Representative images of haematoxylin/eosin stained liver sections from WT and Alb-GDF15$^{KO}$ mice after 24 weeks of HFD feeding

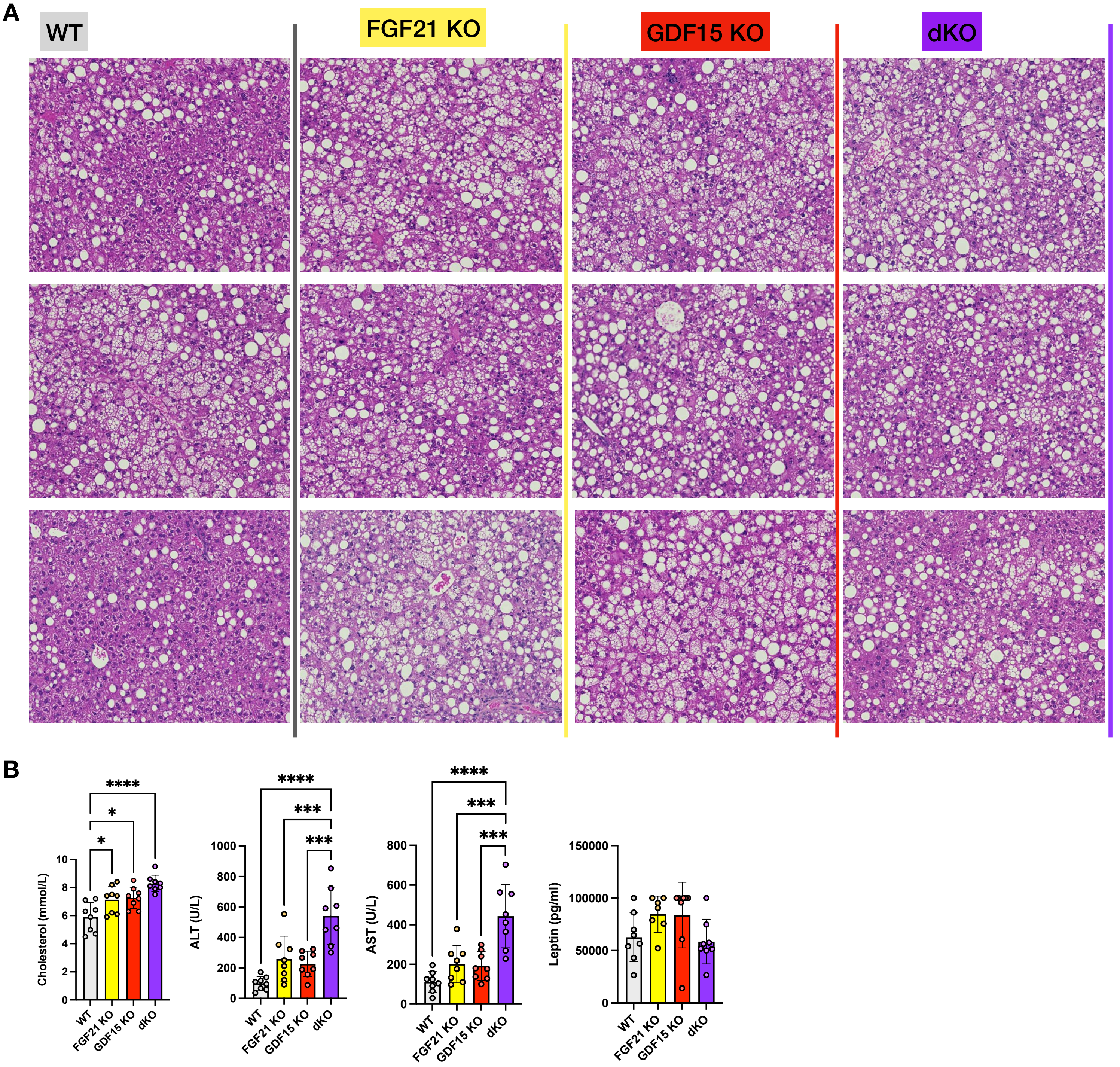


**Figure 6 Supplemental**

(A) Representative images of haematoxylin/eosin stained liver sections from WT, FGF21 KO, GDF15 KO and dKO mice after 25 weeks HFD feeding.

(B) Plasma cholesterol, alanine transaminase (ALT), aspartate transaminase (AST) and leptin from random fed mice, after 24 weeks of HFD-feeding (n=8).
